# Hominoid-specific sulcal variability is related to face perception ability

**DOI:** 10.1101/2022.02.28.482330

**Authors:** Benjamin J. Parker, Willa I. Voorhies, Guo Jiahui, Jacob A. Miller, Ethan Willbrand, Tyler Hallock, Nicholas Furl, Lúcia Garrido, Brad Duchaine, Kevin S. Weiner

**Author notes:** Corresponding author: Kevin S. Weiner.

## Abstract

Human perception requires complex cortical networks that function at neuroanatomical scales of microns and temporal scales of milliseconds. Despite this complexity, what if just one morphological feature of the brain could predict perceptual ability? Here, we tested this hypothesis with pre-registered analyses of neuroanatomy and face perception in neurotypical controls (NTs) and individuals with developmental prosopagnosia (DPs). Results show that the length of the mid-fusiform sulcus (MFS), a hominoid-specific tertiary sulcus in ventral temporal cortex (VTC), was shorter in DPs than NTs. Furthermore, individual differences in MFS length in the right, but not left, hemisphere predicted individual differences in face perception. These results support theories linking brain structure and function to perception, as well as indicate that one feature – variability in MFS length – can predict face perception. Finally, these findings add to growing evidence supporting a role of morphological variability of late developing, tertiary sulci and individual differences in cognition.

## INTRODUCTION

The relationship among brain structure, brain function, and behavior is of major interest in neuroscience, evolutionary biology, and psychology. This relationship is especially intriguing when considering hominoid-specific brain structures because they cannot be examined in widely studied animal models in neuroscience such as mice, marmosets, and macaques (Armstrong et al., 1995; Connolly, 1950; Weiner, 2019). For example, the fusiform gyrus (FG) is a hominoid-specific structure critical for face processing (Duchaine and Yovel, 2015; Kanwisher et al., 1997), object recognition (Gauthier and Tarr, 2016), and reading (Cohen et al., 2000; Wandell et al., 2012). The FG contains a shallow tertiary sulcus, the mid-fusiform sulcus (MFS), that reliably divides the FG into lateral and medial partitions in every hemisphere, and serves as a functional and microarchitectural landmark in humans (Grill-Spector and Weiner, 2014; Weiner et al., 2014; Parvizi et al., 2012). By definition, tertiary sulci such as the MFS are smaller in surface area and shallower in depth compared to primary and secondary sulci, emerge last in gestation, continue to develop after birth, and many are hominoid-specific (Welker, 1990; Sanides, 1964). Immediately relevant for the present study, the MFS exhibits extensive variability across hominoids: it can be as short as a few millimeters or as long as several centimeters in both humans and chimpanzees (Miller et al., 2020; Weiner et al., 2014). However, whether or not this extensive variability reflects individual differences in perception is presently unknown. Further motivating this question are recent findings showing a relationship between either the depth or length of tertiary sulci and cognition in clinical populations and neurotypical controls (NTs). For example, tertiary sulcal depth predicts reasoning skills in children (Voorhies et al., 2021), while tertiary sulcal length is related to whether individuals with schizophrenia will hallucinate or not (Garrison et al., 2015).

Motivated by these findings, we pre-registered analyses (https://osf.io/ydqc4) that leveraged two previously published datasets (Garrido et al., 2009; Jiahui et al., 2018) in which behavioral and anatomical brain data were acquired in 82 participants. A main benefit of these datasets is that they contain data from both NTs, as well as individuals with developmental prosopagnosia (DPs) – individuals who have severe deficits recognizing the faces of familiar people without accompanying insult to the brain (Avidan and Behrmann, 2021; Susilo and Duchaine, 2013). Our pre-registered analyses focused on two main questions: 1) Do morphological features of the MFS differ between DPs and NTs? and 2) Is morphological variability of the MFS predictive of face processing ability in NTs, DPs, or both?

## RESULTS

As outlined in our pre-registration, we first reconstructed the cortical surface for all participants (39 DPs (26 females, mean age 37.1y); 43 NTs (26 females, mean age: 36.7 y)) using FreeSurfer (Dale et al., 1999). We then manually defined the MFS and the two sulci surrounding the FG (occipitotemporal, OTS; collateral, CoS) in each hemisphere (N=492 total sulci; Figure 1A) and implemented a two-pronged analysis approach. First, we compared sulcal morphological features between groups. Second, for those features that were significantly different between groups, we tested if there was a relationship between morphological variability and scores of the Cambridge Face Memory Test (CFMT) within each group. The CFMT is a commonly used measure of unfamiliar face recognition (Duchaine and Nakayama, 2006). Results from all pre-registered anatomical and behavioral analyses are included in the Supplementary Materials.

**Figure 1.**
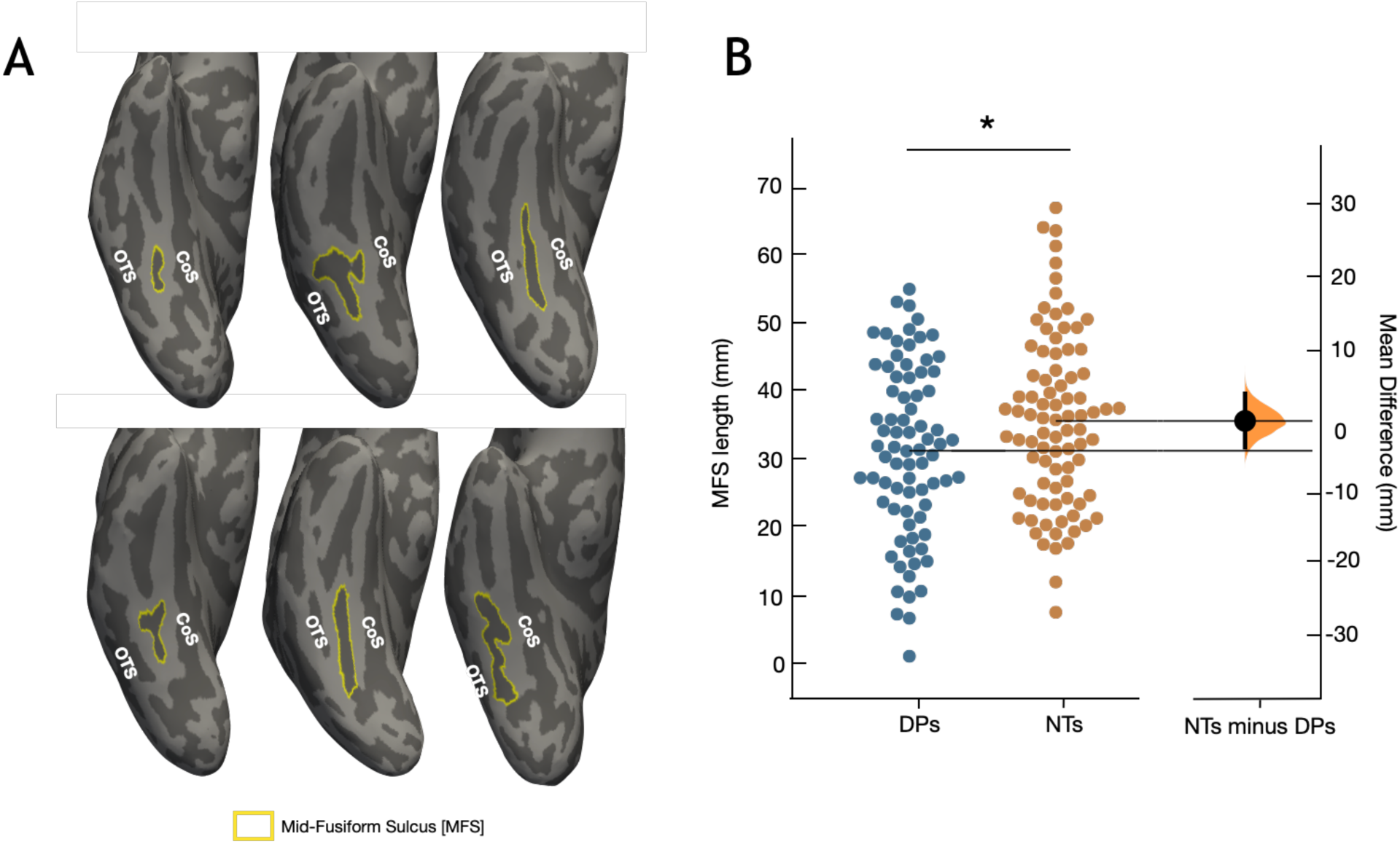
The mid-fusiform sulcus (MFS) is morphologically different between developmental prosopagnosics (DPs) and neurotypical controls (NTs). **A**. Three example inflated cortical surface reconstructions of right hemispheres from DPs (top) and NTs (bottom); the leftmost hemisphere belongs to the 25th percentile of MFS length for its respective group, the center to the 50th percentile, and the rightmost to the 75th percentile. The collateral (CoS) and occipito-temporal (OTS) sulci have been labeled. The mid-fusiform sulcus (MFS) is outlined in yellow in each hemisphere. Dark gray: sulci. Light gray: gyri. **B**. Swarm plot showing MFS length as a function of group (DPs, blue; NTs, orange). On the right y-axis, a bootstrap of 5,000 iterations was used to generate a confidence interval (black bar) displayed with a density plot (orange) which depicts the mean difference in MFS length for each iteration (plot generated with Python package DABEST; Ho et al., 2019). On average in the sample, the MFS is shorter in DPs compared to NTs (*p=.03).

For the first time, we show that MFS length, but not MFS depth, is predictive of face processing in two ways. First, the MFS is shorter in DPs (32.94±12.34) compared to NTs (NTs: 37.36±12.33). A 3-way ANOVA with group, hemisphere, and participant natal sex as factors revealed a main effect of group (F(1, 153)=4.61, p=.03; Figure 1B) and a group x hemisphere x natal sex interaction (F(1, 153)=6.40, p=.02; Supplementary Figure 6). Second, the length of the MFS in the right (r=.34, p=.03), but not left (r=-.11, p=.48), hemisphere predicted CFMT scores in NTs and these correlations were significantly different from one another (Fisher’s Z=2.08, p=.04; Figure 2A). The opposite was true in DPs: the length of the MFS in the left (r=-.34, p=.04), but not right (r=-.14, p=.38), hemisphere predicted CFMT score, but these correlations were not significantly different from each other (Fisher’s Z=.89, p=.38). While this difference between groups and hemispheres may seem surprising, when including all participants as a single distribution with a range of CFMT scores (in which DPs are on the lower end of this distribution) separately in each hemisphere, the length of the MFS still predicts CFMT score in the right (Figure 2B; r =.23, p =.04), but not the left (Figure 2B; r =.06, p =.56), hemisphere. Thus, the positive correlation between MFS length in the right hemisphere and CFMT scores is stronger in NTs, but also reflects a general relationship when considering both groups together. However, the negative correlation between MFS length in the left hemisphere and CFMT scores in DPs does not reflect a general relationship when considering both groups together.

**Figure 2.**
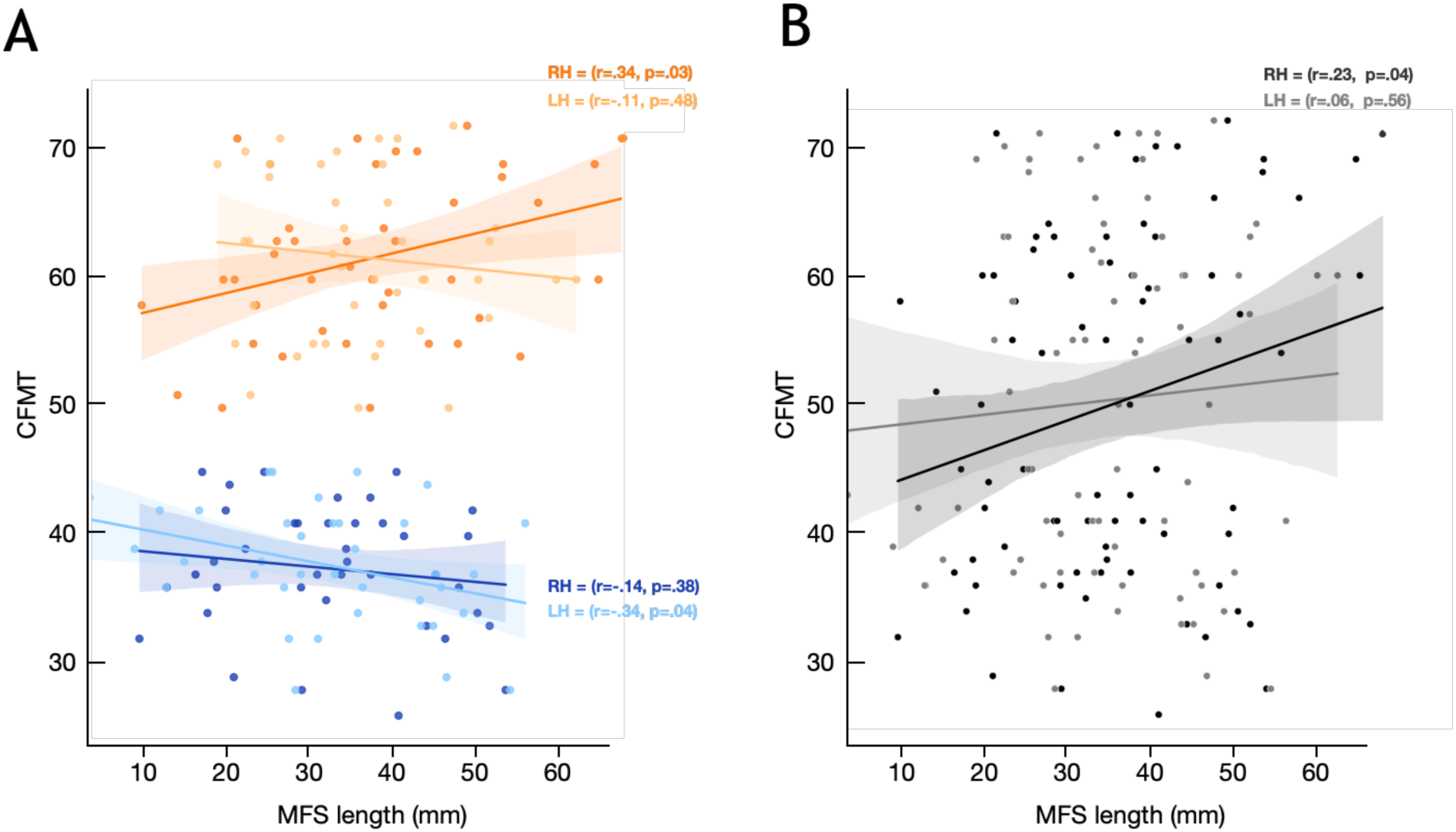
MFS length is correlated with face perception ability. **A**. CFMT performance as a function of MFS length in NTs (orange) and DPs (blue) for the right (darker shade) and left (lighter shade) hemispheres. The length of the MFS in the NT right (r=.34, p=.03), but not NT left (r=.11, p=.48), hemisphere predicted CFMT scores, and these correlations were significantly different from one another (Fisher’s z=2.08, p=.04). In DPs, the length of the MFS in the left (r=-.34, p=.04), but not right (r=-.14, p=.38), hemisphere predicted CFMT score, but these correlations were not significantly different from each other (Fisher’s z=.89, p=.38). **B**. CFMT performance as a function of MFS length collapsed across groups in the left (light gray) and right (black) hemispheres. While it may seem surprising that a longer MFS in the right hemisphere is related to higher CFMT scores in NTs, while the opposite is true in DPs in the left hemisphere, when considering all of the data together as one distribution of participants with a range of CFMT scores (in which DPs are on the lower end of this distribution), there is a positive correlation between MFS length and CFMT score in the right hemisphere (r=.23, p=.04), but not the left hemisphere (r=.06, p=.56).

Finally, while there was no difference in depth between groups for any of the three sulci (F(1, 156)=.60, p=.40), the MFS was shallower in NTs compared to DPs who identified as male, while the opposite was true for those who identified as female (natal sex x group interaction: F(1, 156)=5.10, p=.03); Supplementary Figure 3A). Despite this interaction, MFS depth did not predict CFMT score in either group (NTs: F(1, 76)=2.7, p=.10; DPs: F(1,70)=2.40, p=.12).

## DISCUSSION

Together, the results of our pre-registered analyses reveal that hominoid-specific sulcal variability is related to face perception ability in two main ways. First, the length of the MFS was shorter in DPs than NTs. Second, MFS length in the right hemisphere was positively correlated with performance on a standard measure of face recognition. As the MFS is a hominoid-specific structure, together, these findings empirically support that hominoid-specific sulcal variability is related to face perception for the first time.

These two main findings support a classic anatomical theory (Sanides, 1962, 1964) positing that tertiary sulci are important landmarks in association cortices. They also complement recent findings showing a relationship between either the depth or length of tertiary sulci and cognition in clinical populations and NTs (Voorhies et al., 2021; Yao et al., 2021; Brun et al., 2016; Garrison et al., 2015). Extending these findings, the present results are the first to i) relate individual differences in tertiary sulcal length to individual differences in perception and ii) differentiate DPs from NTs based on tertiary sulcal length.

As the MFS is a functional and anatomical landmark (Weiner, 2019; Grill-Spector and Weiner, 2014), the relationship between MFS length and face perception identified here is likely associated with several functional and anatomical differences across NTs and DPs, which can be explored in future research. Importantly, this structural-functional coupling is not epiphenomenal: electrical charge delivered to electrodes on cortex lateral, but not medial, to the MFS produces causal, face-specific perceptual distortions (Jonas and Rossion, 2021; Parvizi et al., 2012). Thus, future research can quantify the relationship between MFS length and (i) transitions in large-scale functional maps, (ii) the location of fine-scale functional regions, and (iii) decreased category-selectivity in DPs compared to NTs in VTC (Jiahui et al., 2018). Additionally, the MFS also identifies microstructural transitions in VTC based on cytoarchitecture, myelin content, and receptor architecture (Weiner and Yeatman, 2020). As such, the relationship between MFS length and face perception identified here could also be related to differences in the microstructure within and around the MFS in DPs compared to NTs, which is consistent with the influential neural migration theory (Ramus, 2004) proposed to underlie selective developmental disorders such as DP. Furthermore, recent findings show that anatomical features of longitudinal white matter tracts positioned lateral to the MFS are different between DPs and NTs in which these differences correlate with face perception (Gomez et al., 2015). Altogether, the seemingly simple relationship between MFS length and face perception likely reflects complex, multiscale functional and anatomical differences between NTs and DPs.

More broadly, individual differences in MFS morphology may not only be linked to individual differences in face perception and DP, but may also extend to additional disorders. For example, a recent study found that MFS morphology predicts behavioral performance on a theory of mind task in individuals with Autism Spectrum Disorder (Ammons et al., 2021). Additionally, Bouhali and colleagues (2019) recently identified a cortical region selective for graphemes overlapping with the MFS (Bouhali et al., 2019). Thus, the morphology of the MFS may also be a critical biomarker in individuals with dyslexia or even predict phonological skills across individuals without dyslexia, which can be explored in future research. Finally, our study opens the door to shifting the focus from morphological analyses of primary structures in different syndromes and diseases to tertiary structures (such as the MFS) which emerge later in gestation, continue to develop after birth, and are either hominoid- or human-specific, which makes them ideal targets for future studies striving to better understand the neuroanatomical underpinnings of human developmental disorders.

## Acknowledgements

This research was supported by a T32 HWNI training grant (Parker), as well as start-up funds from UC Berkeley (Weiner). Data collected in London was supported by an ESRC grant (RES-061-23-0400) to BD; data collection at Dartmouth was supported by a Rockefeller Foundation award.

## MATERIALS AND METHODS

Here, we include the Materials and Methods as written in our pre-registered analyses (modified to include past tense). We also include our original predictions and rationale as well if we specified them in the pre-registration (https://osf.io/fzb48/).

### Data Acquisition

Our analyses were conducted using two previously published datasets from Garrido et al., 2009 (Dataset 1) and Jiahui et al., 2018 (Dataset 2), which we describe in turn below.

#### Dataset 1 (Garrido et al., 2009)

Participants. Seventeen individuals with developmental prosopagnosia (DPs; 11 females, mean age 30.94 years (SD = 7.54, range 20–46)) and 18 neurotypical controls (NTs; 11 females, 28.94 (SD = 5.70, range 23–43)). All reported being right-handed. All 35 participants showed normal or corrected to normal visual acuity when tested with Test Chart 2000 (Thompson Software Solutions, Hatfield, UK). DP participants contacted the Duchaine lab through http://www.faceblind.org and reported significant difficulties recognizing familiar faces in everyday life. To ascertain that the face recognition deficits that the participants were reporting were consistent with DP, each individual was tested on the Cambridge Face Memory Test (CFMT; Duchaine and Nakayama, 2006) and on a famous faces test. All DPs performed significantly below the mean of published NT means for these two tests.

Brain Data Acquisition. Each participant was scanned on a 3T whole-body MRI scanner (Magnetom TIM Trio, Siemens Medical Systems, Erlangen, Germany) operated with a radio frequency body transmit and 12-channel receive head coil. For each participant, a T1-weighted (T1w), 3D-modified driven equilibrium Fourier transform (MDEFT) dataset was acquired in sagittal orientation with 1 mm isotropic resolution (176 partitions, field of view = 256 × 240 mm^2^, matrix 256 × 240 × 176) with the following parameters: repetition time = 7.92 ms, echo time = 2.48 ms, inversion time = 910 ms (symmetrically distributed around the inversion pulse; quot =50%), flip angle α = 16x, fat saturation, bandwidth 195 Hz/pixel. The sequence was specifically optimized for reduced sensitivity to motion, susceptibility artifacts, and B_1_ field inhomogeneities.

Behavioral tasks. All participants were tested on a battery of tests tapping different aspects of face and object processing. Results on 11 behavioral tasks were published in Garrido et al., 2009, including the CFMT and the Famous Faces Test described above, as well as three Old-New Recognition Tasks (faces, cars, and horses) (Duchaine and Nakayama, 2006), the Cambridge Face Perception Test (Duchaine et al., 2007), the Cambridge Hair Memory Test (Garrido et al., 2009), a Sequential Matching Task for Identity and a Sequential Matching Task for Emotion (Garrido et al., 2009), the Reading the Mind in the Eyes Test (Baron-Cohen et al., 2001), and the Films Facial Expression Task (Garrido et al., 2009).

#### Dataset 2 (Jiahui et al., 2018)

Participants. Twenty-two DPs (seven males, mean age 41.9 y) and 25 NTs (10 males, mean age 42.3 y) participated in the study. DPs were recruited from www.faceblind.org, and all reported problems in daily life with face recognition. To assess their face recognition, DPs were tested with the Cambridge Face Memory Test (CFMT), a famous face test, and an old–new face discrimination test. All but one DP performed two or more standard deviations (SDs) below the mean of published control results in at least two of the three diagnostic tests. The DP participant who did not reach -2 SD on two tests scored poorly on two of the three tasks (CFMT: z = -1.9; famous face: z = -7.1; old– new: z =-0.5), so we included them to increase the sample size. All participants had normal or corrected-to-normal vision and had no current psychiatric disorders. Participants provided written informed consent before doing the tasks, and all procedures were approved by Dartmouth’s Committee for the Protection of Human Participants.

Brain Data Acquisition. All participants were scanned in a 3.0T Philips MR scanner (Philips Medical Systems, WA, USA) with a SENSE (SENSitivity Encoding) 32-channel head coil. A high-resolution anatomical volume was acquired at the beginning of the scan using a high-resolution 3D magnetization-prepared rapid gradient-echo sequence (220 slices, field of view = 240 mm, acquisition matrix = 256 × 256, voxel size = 1 × 0.94 × 0.94 mm).

Face processing tasks. All participants (except one NT) were tested on the CFMT, and DP participants were also tested on a Famous Faces Test (all except two DPs) and on the Old-New Recognition Test for faces (all but one DP).

### Data Analyses

For our analyses, we implemented a two-fold approach. First, we compared morphological features of three sulci in VTC (MFS, CoS, and OTS) between DPs and NTs. Second, we quantified the relationship between morphological features and behavioral performance on face processing tasks. Each approach is explained in turn below.

Morphological analyses. All sulci were defined by BP, EW, TH, and KSW, blind to the identity of all participants. Specifically, from each T1w scan in Datasets 1 and 2, BP, EW, TH, and KSW generated cortical surface visualizations using FreeSurfer (FS). The authors then manually identified the 3 sulci of interest in each hemisphere using protocols from our previously published work (Weiner et al., 2014; Miller et al., 2020). Specifically, a three-tiered approach was implemented to identify the MFS, OTS, and CoS. First, the FG was identified as the major gyrus in VTC. Second, once the FG was identified, the OTS, CoS, and MFS were identified based on the following criteria: 1) the CoS was identified as a deep and long sulcus identifying the medial extent of the FG, 2) the OTS was identified as a deep and long sulcus identifying the lateral extent of the FG, and 3) the MFS was identified as either a single shallow longitudinal sulcus dividing the FG into lateral and medial partitions or it was identified as two or more shallow sulcal components dividing the FG into lateral and medial partitions. In addition to these criteria, the MFS varies morphologically across participants and between hemispheres regarding its intersections with the CoS and OTS, as well as regarding the number of components. Third, as the MFS has predictable anterior (posterior extent of the hippocampus) and posterior (posterior transverse collateral sulcus) landmarks, but the OTS and CoS can extend longitudinally from the occipital pole to the temporal pole, we restricted our OTS and CoS definitions to the portions in VTC surrounding the MFS. This is consistent with our previous protocols and assures that the portions of the OTS and CoS being compared to the MFS are within the VTC and are approximately equal along the length of the three sulci. Thus, this assures that any morphological differences between groups is not due to the fact that the OTS and CoS are much longer than the MFS, for example. Building from previous analyses among the CoS, OTS, and MFS (Weiner et al., 2014) as well as previous anatomical comparisons between DPs and NTs (Garrido et al., 2009; Behrmann et al., 2007), morphological analyses focused on three main anatomical features: (1) sulcal depth, (2) cortical thickness, and (3) sulcal length (just for the MFS since we restricted the length of the OTS and CoS to the portions surrounding the MFS). Using functions in FS and custom software, we measured mean sulcal depth and cortical thickness for each of the 3 sulci. As in our previous work (Miller et al., 2020), mean sulcal depth was normalized in each individual based on the deepest point in cortex. Additionally, cortical thickness was normalized based on the thickest point in cortex. As our previous work showed that sulcal length is the most variable morphological feature of the MFS across both human and non-human hominoid participants (Weiner et al., 2014; Miller et al., 2020), we measured sulcal length of the MFS (in units of millimeters).

Group differences of morphological measures. After manually defining all sulci in both groups, GJ, LG, and BD provided a list of participants’ codes and their group to BP, EW TH, WV, JM and KSW. DPs and NTs from both datasets (N=82) were then included together for morphological analyses. Specifically, groups were included for each of the dependent measures with three main statistical tests outlined below with rationale and predictions:

1) Rationale: From our previous work, sulcal depth of the MFS is the shallowest compared to OTS (2nd shallowest) and CoS (deepest; Weiner et al., 2014; Miller et al., 2020). As such, we conducted an N-way analysis of variance (ANOVA) on sulcal depth, with sulcus (CoS, MFS, OTS), hemisphere (RH, LH), natal sex (male, female) and group (DP, NTs) as factors.

Prediction: The MFS depth should be shallowest, CoS should be deepest, and OTS should be in between for both groups. Additionally, we hypothesized that a) there will not be group differences in sulcal depth between DPs and NTs for the CoS and OTS given previous findings (Behrmann et al., 2007), and b) there will be group differences in sulcal depth for the MFS, but we do not make any predictions regarding the directionality of the differences.

2) Rationale: From our previous work (Garrido et al., 2009), voxel-based morphometry (VBM) analyses showed differences in the Fusiform Gyrus between DPs and NTs. As grey matter volume as measured with VBM may also be related to cortical thickness, we plan to conduct a 3-way ANOVA on cortical thickness, with sulcus (CoS, MFS, OTS), hemisphere (RH, LH), natal sex (male, female) and group (DP, NTs) as factors.

Prediction: As our previous findings showed that DPs had reduced gray matter volume in the middle FG compared to NTs (Garrido et al., 2009) and portions of each of these three sulci were likely included in the region identified in this previous work, we hypothesized that there will be group differences in cortical thickness for all or a subset of these three sulci. Specifically, we hypothesized that it may be the case that each sulcus is thinner in DPs compared to NTs, which would be consistent with reduced gray matter volume, but we cannot be sure as the region identified previously also contained gyral components. Thus, we did not make an explicit prediction regarding directionality, but did predict that there will be differences in cortical thickness of all or a subset of these three sulci between the two groups.

3) Rationale. While the shallowness of the MFS is its most stable feature, the length of the MFS is its most variable feature: it can be as short as just under 3mm or as long as 5.5 cm (Weiner et al., 2014; Miller et al., 2020). Thus, we measured the length of the MFS in both DPs and NTs, and then compared the length of the MFS between DPs and NTs in both hemispheres.

Prediction: Based on previous studies showing that the length of the MFS is its most variable feature (Weiner et al., 2014; Weiner, 2019; Miller et al., 2020), it may be the case that the MFS will be equally variable between DPs and NTs. Nevertheless, given the large variability in MFS length, it may also be the case that we find group differences in the length of the MFS. As such, we do not explicitly predict one or the other as both outcomes are possible.

## EXPLORATORY ANALYSES

Examining the relationship between brain and behavior. We aimed to perform exploratory analyses examining the relationship between morphological features of VTC sulci and behavior in our two datasets. For any anatomical features that either showed a main effect of group or were included in any interactions from our ANOVA analyses, we performed correlation analyses to explore the relationship between these anatomical features and behavioral performance, including age and natal sex as covariates. We started with the CFMT as a behavioral measure, as that is the only measure that we have for all participants. We then repeated all analyses within each dataset with the behavioral measures available for each dataset.

Relationship between morphological measures. Our recent work (Miller et al., 2020) showed a relationship between cortical thickness and depth particularly with the MFS in a group of NTs from the Human Connectome Project. Concomitantly, we calculated the correlation between thickness and depth for each of the three sulci and for each of the two groups. Based on these previous findings, we predicted that there would be a relationship between thickness and depth for the MFS in our cohort of subjects. However, we did not make an explicit prediction regarding if this relationship would a) occur in one group, but not the other, or b) would be stronger in one group compared to another. Thus, this analysis was also exploratory in nature.

