## Supplementary Materials for "Hominoid-specific sulcal variability is related to face perception ability"

**Parker et al.**

**Supplementary Results**

EXPLORATORY ANALYSES

In our pre-registration, it was stated that exploratory analyses would include all analyses of all behavioral measurements for each dataset. The following section enumerates the results of these tests, and is separated by dataset. In Dataset 1 (Garrido et. al, 2009), the following behavioral metrics were gathered beyond the CFMT reported in the main text: Famous Faces Test (Duchaine and Nakayama, 2005), Old-New Recognition (Duchaine and Nakayama, 2005): faces, cars and horses, Cambridge Face Perception Test – CFPT (Duchaine et al., 2007a; Duchaine et al., 2007b), Sequential Matching of Facial Identity and Facial Expression (Banissy et al., 2011), Cambridge Hair Memory Test - CHMT (Duchaine and Nakayama, 2006), 'Reading the Mind in the Eyes' Test (Baron-Cohen et al. 2001), Films Facial Expression Task (Avidan et al., 2011), and the Weschler Abbreviated Intelligence Scale (WASII; full, performance, and verbal). In addition to raw scores on these tests, we also include reaction time where appropriate.

All ANOVAs were constructed to predict behavioral score with factors of participant gender (*male, female*), participant group (*DP, NT*), and MFS sulcal length (mm). After correcting alpha with Bonferroni's post-hoc adjustment ($p<.002$), none of the models predicting behavior with MFS length as a factor were significant. Note that this was the case when the tests were conducted collapsed across groups (**Supplementary Fig. 2.1)**, and when analyzing DPs and NTs separately (**Supplementary Fig. 2.2 and 2.3)**.

In addition to MFS sulcal length, we also pre-registered 3-way ANOVAs with cortical thickness and sulcal depth respectively, each with sulcus (*CoS, MFS, OTS*), hemisphere (*RH, LH*), gender (*male, female*) and group (*DP, NT*) as factors. While there is no main effect of group ($F(1, 468)<2.0$, $p>.15$) (**Supplementary Fig. 3bi-iii)**, there is a significant group x participant gender interaction in which NT males tend to have cortically thicker VTC sulci than their DP counterparts, while NT females tend to have cortically thinner VTC sulci than their DP counterparts ($F(1,468)=10.0$, $p<.002$) (**Supplementary Fig. 3ai-iii)**.

As this interaction closely mirrored the interaction for MFS sulcal depth, we examined the relationship between sulcal depth and cortical thickness using Pearson's correlation separately for each sulcus, hemisphere, and group (**Supplementary Fig. 4)**. After correcting with Bonferroni's post-hoc adjustment ($p<.004$), only the NT right hemisphere ($r=-.51$, $p<.0005$) and the DP left hemisphere ($r=-.48$, $p<.004$) were significantly anticorrelated for the MFS, but not OTS and CoS.

Dataset 1 (Garrido et al., 2009)

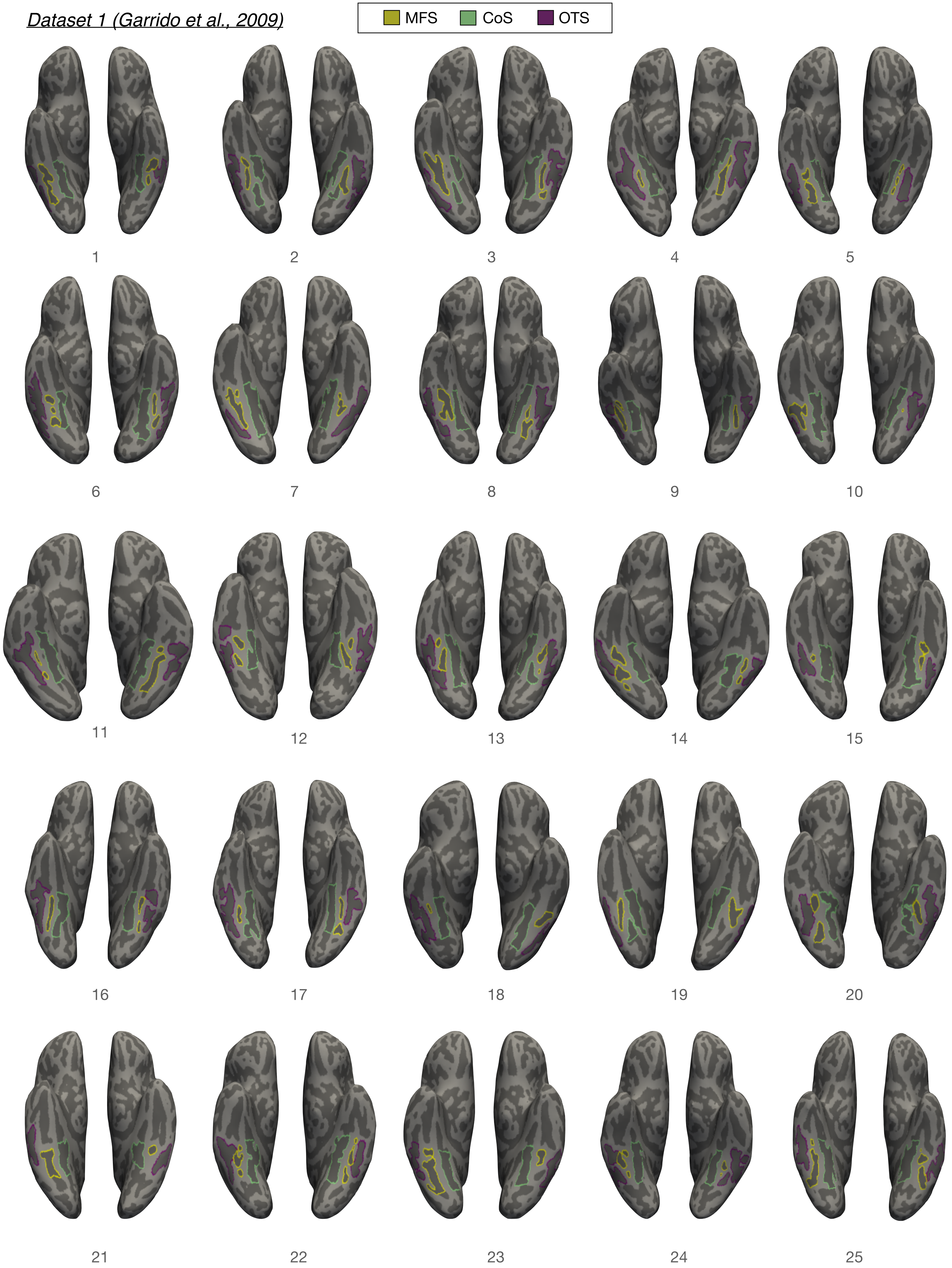

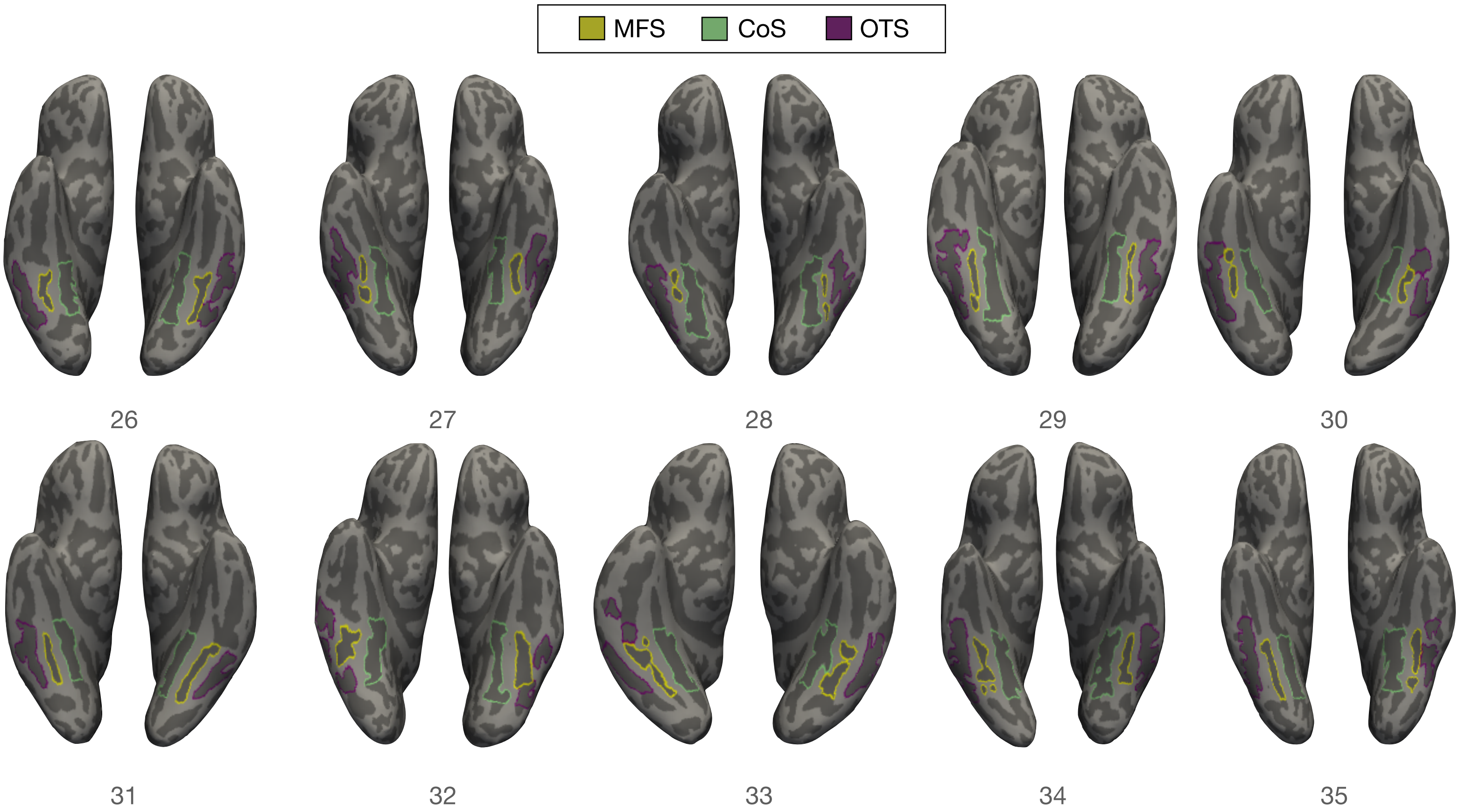
MFS
CoS
OTS

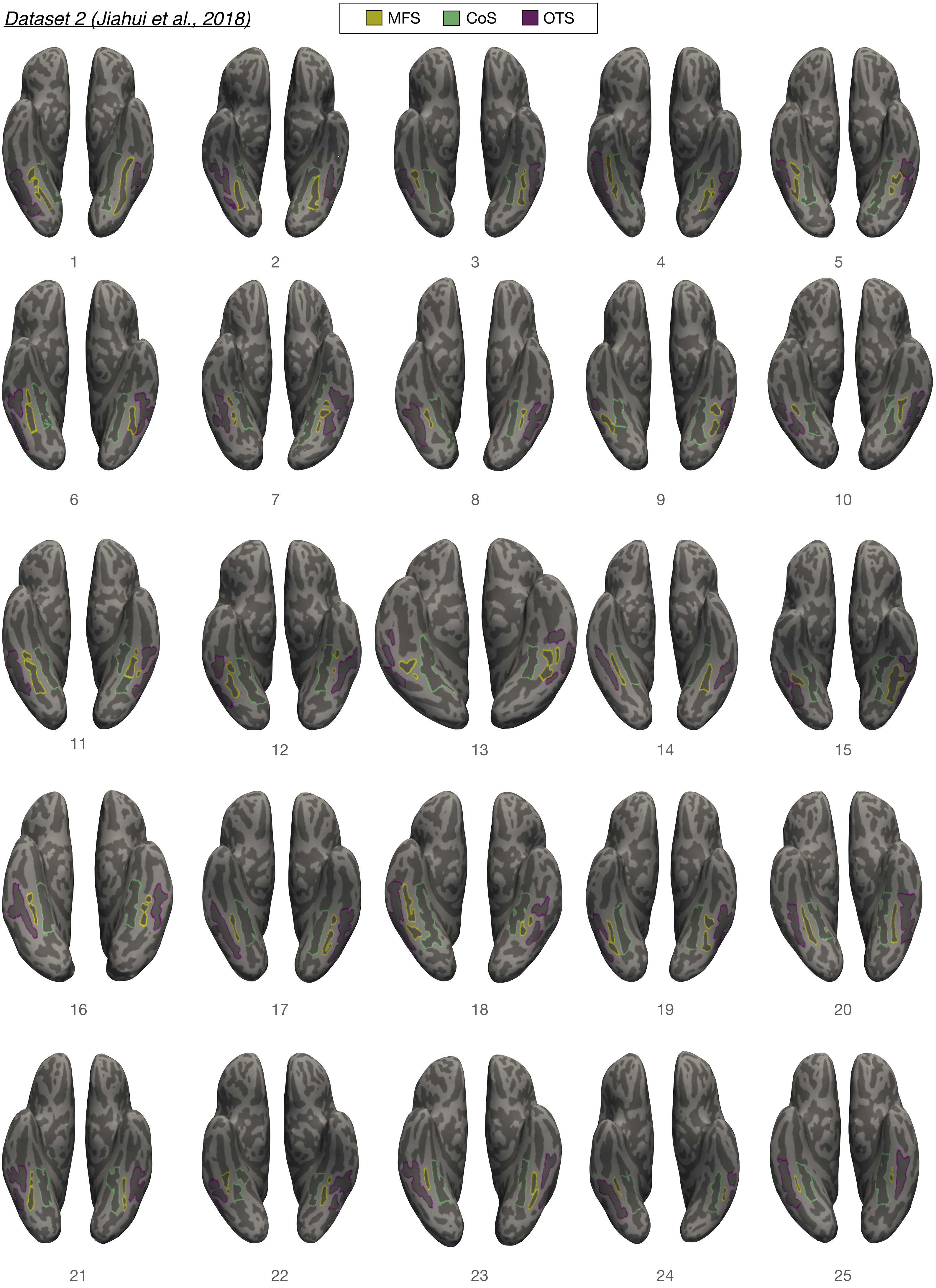

Dataset 2 (Jiahui et al., 2018)
MFS
CoS
OTS

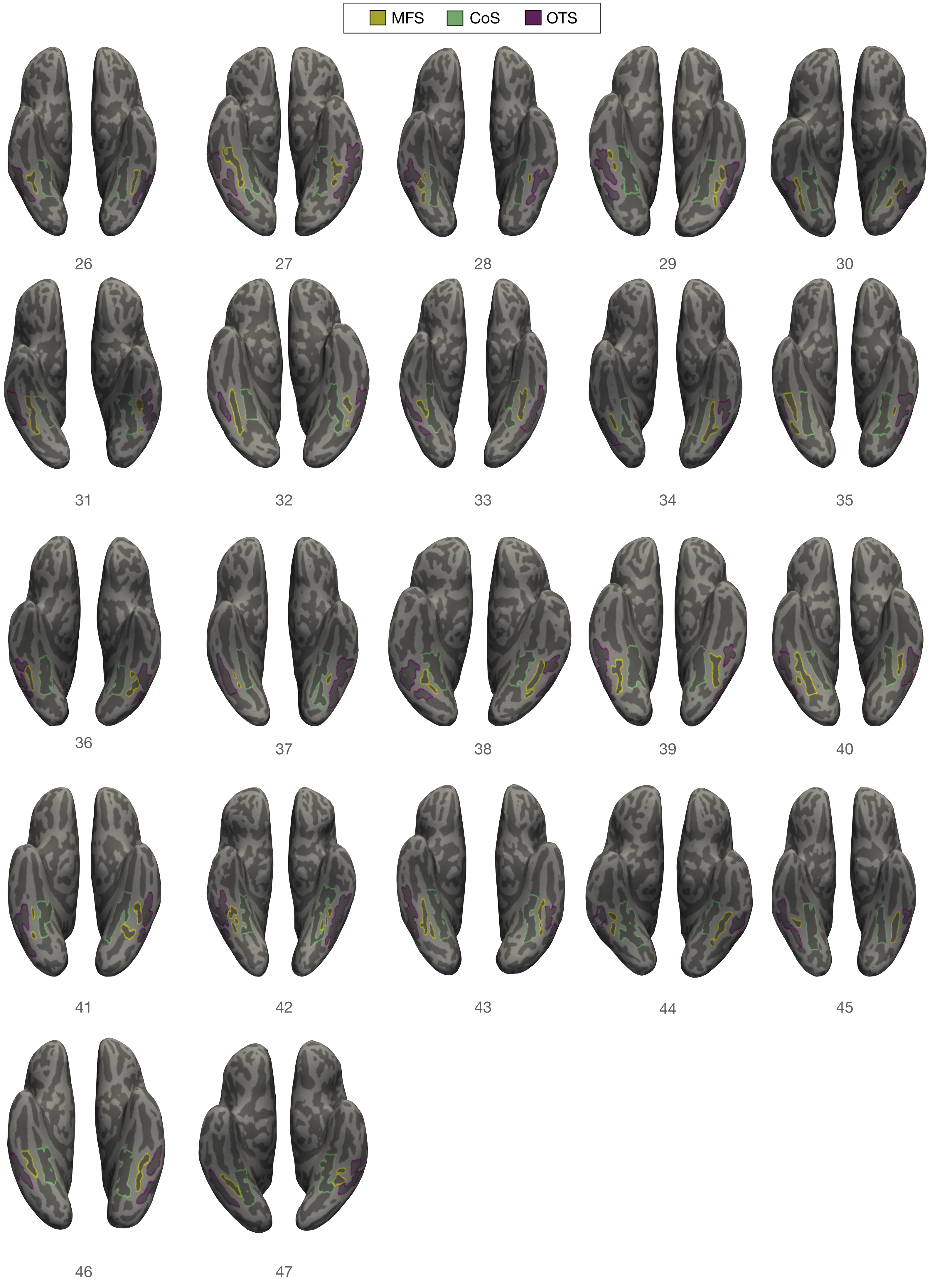
MFS
CoS
OTS

**Supplementary Figure 1. VTC sulci in every individual participant from Datasets 1 and 2.** Manual sulcal labels in the left and right hemispheres of each participant. Each sulcus is displayed on the inflated cortical surface in FreeSurfer 6.0.0 and is colored according to the key at the top: OTS (yellow), MFS (Blue), CoS (Green). Only those portions of the CoS and OTS within VTC surrounding the MFS were defined (Materials and Methods).

**a.** Baldwomen

| factors | sum_sq | df | F | PR(>F) |
|---|---|---|---|---|
| C(Sex) | 0.010724 | 1 | 1.122262 | 0.293609 |
| max_path_length | 0.002558 | 1 | 0.267711 | 0.606742 |
| max_path_length:C(Sex) | 0.001677 | 1 | 0.175539 | 0.676707 |
| Age | 0.004045 | 1 | 0.423279 | 0.517749 |
| Age:C(Sex) | 0.000166 | 1 | 0.017394 | 0.89551 |
| max_path_length:Age | 0.015576 | 1 | 1.630045 | 0.206536 |
| max_path_length:Age:C(Sex) | 0.003662 | 1 | 0.383198 | 0.538202 |
| Residual | 0.582884 | 61 | NaN | NaN |

**b.** Baldwomen RT

| factors | sum_sq | df | F | PR(>F) |
|---|---|---|---|---|
| C(Sex) | 4690568.26647 | 1 | 6.766658 | 0.011639 |
| max_path_length | 119094.217495 | 1 | 0.171806 | 0.679965 |
| max_path_length:C(Sex) | 118263.233621 | 1 | 0.170608 | 0.681021 |
| Age | 3128592.889047 | 1 | 4.513338 | 0.037693 |
| Age:C(Sex) | 5561351.638573 | 1 | 8.022859 | 0.006251 |
| max_path_length:Age | 51681.596588 | 1 | 0.074556 | 0.785737 |
| max_path_length:Age:C(Sex) | 306901.801036 | 1 | 0.442739 | 0.508311 |
| Residual | 42284484.719406 | 61 | NaN | NaN |

**c.** CFPT INV

| factors | sum_sq | df | F | PR(>F) |
|---|---|---|---|---|
| C(Sex) | 1042.605152 | 1 | 4.637664 | 0.035239 |
| max_path_length | 607.455392 | 1 | 2.702053 | 0.105364 |
| max_path_length:C(Sex) | 189.665781 | 1 | 0.843662 | 0.361969 |
| Age | 221.861093 | 1 | 0.986871 | 0.324433 |
| Age:C(Sex) | 3.839552 | 1 | 0.017079 | 0.896454 |
| max_path_length:Age | 2.57875 | 1 | 0.011471 | 0.91506 |
| max_path_length:Age:C(Sex) | 290.286646 | 1 | 1.291238 | 0.260264 |
| Residual | 13713.567024 | 61 | NaN | NaN |

**d.** CFPT UP

| factors | sum_sq | df | F | PR(>F) |
|---|---|---|---|---|
| C(Sex) | 1.896769 | 1 | 0.007229 | 0.932523 |
| max_path_length | 643.707948 | 1 | 2.453166 | 0.122461 |
| max_path_length:C(Sex) | 11.173566 | 1 | 0.042582 | 0.837201 |
| Age | 226.326842 | 1 | 0.86253 | 0.356692 |
| Age:C(Sex) | 374.149767 | 1 | 1.425882 | 0.237061 |
| max_path_length:Age | 22.328159 | 1 | 0.085092 | 0.771501 |
| max_path_length:Age:C(Sex) | 35.783074 | 1 | 0.136369 | 0.713197 |
| Residual | 16006.327278 | 61 | NaN | NaN |

**c.** CHMT

| factors | sum_sq | df | F | PR(>F) |
|---|---|---|---|---|
| C(Sex) | 61.598923 | 1 | 2.168827 | 0.145975 |
| max_path_length | 10.00594 | 1 | 0.352298 | 0.55501 |
| max_path_length:C(Sex) | 0.270481 | 1 | 0.009523 | 0.92258 |
| Age | 2.660828 | 1 | 0.093685 | 0.760586 |
| Age:C(Sex) | 20.997861 | 1 | 0.73931 | 0.393249 |
| max_path_length:Age | 2.152193 | 1 | 0.075776 | 0.784035 |
| max_path_length:Age:C(Sex) | 8.286033 | 1 | 0.291742 | 0.591073 |
| Residual | 1732.519365 | 61 | NaN | NaN |

**d.** Eyes Test emotion

| factors | sum_sq | df | F | PR(>F) |
|---|---|---|---|---|
| C(Sex) | 0.002598 | 1 | 0.000304 | 0.986144 |
| max_path_length | 0.042783 | 1 | 0.005008 | 0.943812 |
| max_path_length:C(Sex) | 0.021165 | 1 | 0.002478 | 0.960463 |
| Age | 7.58409 | 1 | 0.887837 | 0.349781 |
| Age:C(Sex) | 42.123875 | 1 | 4.93126 | 0.030099 |
| max_path_length:Age | 29.167925 | 1 | 3.414563 | 0.069473 |
| max_path_length:Age:C(Sex) | 69.439175 | 1 | 8.128945 | 0.005937 |
| Residual | 521.074986 | 61 | NaN | NaN |

**e.** Famous Faces UK Exposed

| factors | sum_sq | df | F | PR(>F) |
|---|---|---|---|---|
| C(Sex) | 7.258304 | 1 | 0.326462 | 0.569849 |
| max_path_length | 5.990166 | 1 | 0.269424 | 0.605597 |
| max_path_length:C(Sex) | 49.408519 | 1 | 2.222283 | 0.141185 |
| Age | 8.671491 | 1 | 0.390024 | 0.534616 |
| Age:C(Sex) | 1.55349 | 1 | 0.069872 | 0.792414 |
| max_path_length:Age | 47.288511 | 1 | 2.12693 | 0.14986 |
| max_path_length:Age:C(Sex) | 29.426754 | 1 | 1.323548 | 0.254446 |
| Residual | 1356.226893 | 61 | NaN | NaN |

**f.** Famous Faces UK Identified

| factors | sum_sq | df | F | PR(>F) |
|---|---|---|---|---|
| C(Sex) | 4.507592 | 1 | 0.019274 | 0.890042 |
| max_path_length | 22.465201 | 1 | 0.096057 | 0.75767 |
| max_path_length:C(Sex) | 267.747826 | 1 | 1.144836 | 0.288848 |
| Age | 3.232419 | 1 | 0.013821 | 0.9068 |
| Age:C(Sex) | 357.278923 | 1 | 1.527653 | 0.221203 |
| max_path_length:Age | 246.734979 | 1 | 1.054989 | 0.308417 |
| max_path_length:Age:C(Sex) | 311.815379 | 1 | 1.33326 | 0.252729 |
| Residual | 14266.33673 | 61 | NaN | NaN |

**g.** Films Task Up

| factors | sum_sq | df | F | PR(>F) |
|---|---|---|---|---|
| C(Sex) | 129.671232 | 1 | 3.141702 | 0.081307 |
| max_path_length | 7.171793 | 1 | 0.17376 | 0.678255 |
| max_path_length:C(Sex) | 18.937439 | 1 | 0.45882 | 0.500737 |
| Age | 1.9637 | 1 | 0.047577 | 0.828063 |
| Age:C(Sex) | 38.041983 | 1 | 0.921689 | 0.340822 |
| max_path_length:Age | 14.073568 | 1 | 0.340977 | 0.561419 |
| max_path_length:Age:C(Sex) | 111.769577 | 1 | 2.707977 | 0.104991 |
| Residual | 2517.726494 | 61 | NaN | NaN |

**h.** WASI Full

| factors | sum_sq | df | F | PR(>F) |
|---|---|---|---|---|
| C(Sex) | 5.151921 | 1 | 0.074202 | 0.786298 |
| max_path_length | 306.519715 | 1 | 4.414764 | 0.040062 |
| max_path_length:C(Sex) | 147.652673 | 1 | 2.126623 | 0.150248 |
| Age | 4.747522 | 1 | 0.068378 | 0.794656 |
| Age:C(Sex) | 69.611749 | 1 | 1.002609 | 0.320912 |
| max_path_length:Age | 0.010571 | 1 | 0.000152 | 0.990198 |
| max_path_length:Age:C(Sex) | 0.632263 | 1 | 0.009106 | 0.92431 |
| Residual | 3957.543908 | 57 | NaN | NaN |

**i.** WASI Verbal

| factors | sum_sq | df | F | PR(>F) |
|---|---|---|---|---|
| C(Sex) | 83.019659 | 1 | 0.65095 | 0.423127 |
| max_path_length | 437.318438 | 1 | 3.428975 | 0.069244 |
| max_path_length:C(Sex) | 179.353879 | 1 | 1.406298 | 0.240592 |
| Age | 44.812578 | 1 | 0.351371 | 0.555682 |
| Age:C(Sex) | 114.438301 | 1 | 0.897301 | 0.347505 |
| max_path_length:Age | 16.093462 | 1 | 0.126187 | 0.723729 |
| max_path_length:Age:C(Sex) | 48.576416 | 1 | 0.380883 | 0.539588 |
| Residual | 7269.563349 | 57 | NaN | NaN |

**j.** WASI Performance

| factors | sum_sq | df | F | PR(>F) |
|---|---|---|---|---|
| C(Sex) | 41.167959 | 1 | 0.8654 | 0.356155 |
| max_path_length | 73.311652 | 1 | 1.5411 | 0.219539 |
| max_path_length:C(Sex) | 41.67519 | 1 | 0.876063 | 0.353231 |
| Age | 3.251898 | 1 | 0.068359 | 0.794684 |
| Age:C(Sex) | 16.508214 | 1 | 0.347023 | 0.558131 |
| max_path_length:Age | 19.553487 | 1 | 0.411038 | 0.524013 |
| max_path_length:Age:C(Sex) | 74.527869 | 1 | 1.566666 | 0.215804 |
| Residual | 2711.547073 | 57 | NaN | NaN |

### L. Horses

| factors | sum_sq | df | F | PR(>F) |
|---|---|---|---|---|
| C(Sex) | 0.000002 | 1 | 0.00055 | 0.981363 |
| max_path_length | 0.002237 | 1 | 0.716218 | 0.400695 |
| max_path_length:C(Sex) | 0.000056 | 1 | 0.01808 | 0.89348 |
| Age | 0.000994 | 1 | 0.318219 | 0.574748 |
| Age:C(Sex) | 0.00163 | 1 | 0.521896 | 0.472795 |
| max_path_length:Age | 0.000124 | 1 | 0.039734 | 0.842665 |
| max_path_length:Age:C(Sex) | 0.003653 | 1 | 1.169507 | 0.283759 |
| Residual | 0.19052 | 61 | NaN | NaN |

### m. Horses RT

| factors | sum_sq | df | F | PR(>F) |
|---|---|---|---|---|
| C(Sex) | 2103153.733192 | 1 | 21.255003 | 0.000021 |
| max_path_length | 26650.660643 | 1 | 0.269338 | 0.605654 |
| max_path_length:C(Sex) | 42597.843077 | 1 | 0.430505 | 0.514209 |
| Age | 351060.850963 | 1 | 3.54791 | 0.06439 |
| Age:C(Sex) | 1458678.508508 | 1 | 14.741773 | 0.000296 |
| max_path_length:Age | 16530.926003 | 1 | 0.167066 | 0.684164 |
| max_path_length:Age:C(Sex) | 24490.878701 | 1 | 0.247511 | 0.620622 |
| Residual | 6035867.425433 | 61 | NaN | NaN |

### n. Sequential Em Inv

| factors | sum_sq | df | F | PR(>F) |
|---|---|---|---|---|
| C(Sex) | 15.522569 | 1 | 0.303565 | 0.583669 |
| max_path_length | 39.36114 | 1 | 0.76976 | 0.383734 |
| max_path_length:C(Sex) | 27.066535 | 1 | 0.529323 | 0.469675 |
| Age | 10.090482 | 1 | 0.197333 | 0.658454 |
| Age:C(Sex) | 230.603696 | 1 | 4.509766 | 0.037766 |
| max_path_length:Age | 14.253042 | 1 | 0.278737 | 0.599445 |
| max_path_length:Age:C(Sex) | 62.846399 | 1 | 1.229046 | 0.271945 |
| Residual | 3119.191888 | 61 | NaN | NaN |

### o. Sequential Em emup

| factors | sum_sq | df | F | PR(>F) |
|---|---|---|---|---|
| C(Sex) | 1.341328 | 1 | 0.078805 | 0.779873 |
| max_path_length | 49.399292 | 1 | 2.902292 | 0.093544 |
| max_path_length:C(Sex) | 77.621033 | 1 | 4.560367 | 0.036743 |
| Age | 113.515131 | 1 | 6.669206 | 0.012225 |
| Age:C(Sex) | 20.538923 | 1 | 1.206696 | 0.276304 |
| max_path_length:Age | 3.30894 | 1 | 0.194406 | 0.660833 |
| max_path_length:Age:C(Sex) | 16.591282 | 1 | 0.974766 | 0.327395 |
| Residual | 1038.26805 | 61 | NaN | NaN |

### p. Sequential ID Inv

| factors | sum_sq | df | F | PR(>F) |
|---|---|---|---|---|
| C(Sex) | 1.991262 | 1 | 0.134641 | 0.714937 |
| max_path_length | 33.742267 | 1 | 2.281518 | 0.136087 |
| max_path_length:C(Sex) | 4.146049 | 1 | 0.280339 | 0.598401 |
| Age | 452.107811 | 1 | 30.569735 | 0.000001 |
| Age:C(Sex) | 2.478255 | 1 | 0.16757 | 0.683715 |
| max_path_length:Age | 0.307478 | 1 | 0.02079 | 0.885827 |
| max_path_length:Age:C(Sex) | 4.129281 | 1 | 0.279206 | 0.59914 |
| Residual | 902.152934 | 61 | NaN | NaN |

### q. Sequential ID up

| factors | sum_sq | df | F | PR(>F) |
|---|---|---|---|---|
| C(Sex) | 33.164825 | 1 | 0.780424 | 0.38048 |
| max_path_length | 43.052861 | 1 | 1.013107 | 0.318136 |
| max_path_length:C(Sex) | 86.855672 | 1 | 2.043861 | 0.157923 |
| Age | 228.639155 | 1 | 5.380266 | 0.02373 |
| Age:C(Sex) | 2.568925 | 1 | 0.060451 | 0.80661 |
| max_path_length:Age | 25.554116 | 1 | 0.601332 | 0.441066 |
| max_path_length:Age:C(Sex) | 1.150257 | 1 | 0.027067 | 0.869864 |
| Residual | 2592.248729 | 61 | NaN | NaN |

## r. PC1

| factors | sum_sq | df | F | PR(>F) |
|---|---|---|---|---|
| C(Sex) | 0.800175 | 1 | 0.942527 | 0.335463 |
| max_path_length | 0.118163 | 1 | 0.139185 | 0.710386 |
| max_path_length:C(Sex) | 0.422453 | 1 | 0.497608 | 0.483238 |
| Age | 0.435364 | 1 | 0.512815 | 0.476655 |
| Age:C(Sex) | 2.240101 | 1 | 2.638617 | 0.109451 |
| max_path_length:Age | 0.540961 | 1 | 0.637199 | 0.427824 |
| max_path_length:Age:C(Sex) | 0.116734 | 1 | 0.137501 | 0.712063 |
| Residual | 51.787028 | 61 | NaN | NaN |

## s. PC2

| factors | sum_sq | df | F | PR(>F) |
|---|---|---|---|---|
| C(Sex) | 0.020632 | 1 | 0.021215 | 0.884675 |
| max_path_length | 1.858138 | 1 | 1.910614 | 0.171936 |
| max_path_length:C(Sex) | 0.71095 | 1 | 0.731028 | 0.395896 |
| Age | 1.08076 | 1 | 1.111281 | 0.295961 |
| Age:C(Sex) | 0.00378 | 1 | 0.003887 | 0.95049 |
| max_path_length:Age | 0.003426 | 1 | 0.003522 | 0.952869 |
| max_path_length:Age:C(Sex) | 1.20836 | 1 | 1.242486 | 0.269365 |
| Residual | 59.32462 | 61 | NaN | NaN |

## t. PC3

| factors | sum_sq | df | F | PR(>F) |
|---|---|---|---|---|
| C(Sex) | 2.285479 | 1 | 2.522364 | 0.117413 |
| max_path_length | 0.139469 | 1 | 0.153924 | 0.696181 |
| max_path_length:C(Sex) | 0.695591 | 1 | 0.767687 | 0.384371 |
| Age | 0.18565 | 1 | 0.204892 | 0.652407 |
| Age:C(Sex) | 5.084795 | 1 | 5.611823 | 0.021022 |
| max_path_length:Age | 0.873925 | 1 | 0.964505 | 0.329934 |
| max_path_length:Age:C(Sex) | 3.503098 | 1 | 3.866187 | 0.053823 |
| Residual | 55.271251 | 61 | NaN | NaN |

### u. Cars RT

| factors | sum_sq | df | F | PR(>F) |
|---|---|---|---|---|
| C(Sex) | 284599.10469 | 1 | 2.683778 | 0.106524 |
| max_path_length | 478.887518 | 1 | 0.004516 | 0.946642 |
| max_path_length:C(Sex) | 100749.212065 | 1 | 0.950068 | 0.333552 |
| Age | 530412.499778 | 1 | 5.001805 | 0.028988 |
| Age:C(Sex) | 1071406.95468 | 1 | 10.103398 | 0.002325 |
| max_path_length:Age | 470786.949195 | 1 | 4.439534 | 0.039236 |
| max_path_length:Age:C(Sex) | 336502.238211 | 1 | 3.173226 | 0.079833 |
| Residual | 6468697.685939 | 61 | NaN | NaN |

### v. Cars

| factors | sum_sq | df | F | PR(>F) |
|---|---|---|---|---|
| C(Sex) | 0.017706 | 1 | 4.182333 | 0.045168 |
| max_path_length | 0.001171 | 1 | 0.276576 | 0.600861 |
| max_path_length:C(Sex) | 0.002044 | 1 | 0.482906 | 0.489748 |
| Age | 0.030308 | 1 | 7.158778 | 0.009565 |
| Age:C(Sex) | 0.001377 | 1 | 0.325191 | 0.570599 |
| max_path_length:Age | 0.000006 | 1 | 0.001356 | 0.970749 |
| max_path_length:Age:C(Sex) | 0.006341 | 1 | 1.497814 | 0.225714 |
| Residual | 0.258252 | 61 | NaN | NaN |

**Supplementary Figure 2.1 Testing for a relationship between MFS length and other behavioral tests in Dataset 1 collapsed across groups: No significant effects.** The following behavioral metrics were gathered beyond the CFMT reported in the main text: [Baldwomen, Baldwomen RT] Old-New Recognition (Duchaine and Nakayama, 2005): faces, [CFPT] Cambridge Face Perception Test (Duchaine et al., 2007a; Duchaine et al., 2007b), [CHMT] Cambridge Hair Memory Test (Duchaine and Nakayama, 2006), [Eyes test emotion] 'Reading the Mind in the Eyes' Test (Baron-Cohen et al. 2001), [Famous faces UK exposed, Famous faces UK exposed] Famous Faces Test (Duchaine and Nakayama, 2005), [Films Task Up] Films Facial Expression Task (Banissy et al, 2011), [WASI Vull, WASI Verbal, WASI Performance] the Weschler Abbreviated Intelligence Scale, [Horses, Horses RT] Old-New Recognition (Duchaine and Nakayama, 2005): horses, [Sequential Em Up, Sequential Em Inv, Sequential ID Inv, Sequential ID Up] Sequential Matching of Facial Identity and Facial Expression (Avidan et al, 2011), [PC1, PC2, PC3] 3 Principal Components identified in Garrido et al. (2009), and [Cars, Cars RT] Old-New Recognition (Duchaine and Nakayama, 2005): cars. The relationship between MFS length and behavior was specific to CFMT performance.

**a.** Baldwomen

| factors | sum_sq | df | F | PR(>F) |
|---|---|---|---|---|
| C(Sex) | 0.009666 | 1 | 1.565269 | 0.222477 |
| max_path_length | 0.003642 | 1 | 0.589777 | 0.449699 |
| max_path_length:C(Sex) | 0.000591 | 1 | 0.095642 | 0.759686 |
| Age | 0.024335 | 1 | 3.940572 | 0.05821 |
| Age:C(Sex) | 0.002265 | 1 | 0.366849 | 0.550186 |
| max_path_length:Age | 0.00599 | 1 | 0.969894 | 0.334143 |
| max_path_length:Age:C(Sex) | 0.000008 | 1 | 0.001247 | 0.972108 |
| Residual | 0.154388 | 25 | NaN | NaN |

**b.** Baldwomen RT

| factors | sum_sq | df | F | PR(>F) |
|---|---|---|---|---|
| C(Sex) | 4835940.293478 | 1 | 4.573806 | 0.04242 |
| max_path_length | 657756.999117 | 1 | 0.622103 | 0.437683 |
| max_path_length:C(Sex) | 1435385.512879 | 1 | 1.35758 | 0.254955 |
| Age | 2459841.105729 | 1 | 2.326504 | 0.13974 |
| Age:C(Sex) | 2814958.914052 | 1 | 2.662373 | 0.115282 |
| max_path_length:Age | 20709.005991 | 1 | 0.019586 | 0.88982 |
| max_path_length:Age:C(Sex) | 402603.896694 | 1 | 0.380781 | 0.542766 |
| Residual | 26432800.338722 | 25 | NaN | NaN |

**c.** CFPT INV

| factors | sum_sq | df | F | PR(>F) |
|---|---|---|---|---|
| C(Sex) | 211.890661 | 1 | 2.215727 | 0.149118 |
| max_path_length | 50.188401 | 1 | 0.524817 | 0.47552 |
| max_path_length:C(Sex) | 41.812543 | 1 | 0.437231 | 0.51451 |
| Age | 188.46409 | 1 | 1.970757 | 0.172668 |
| Age:C(Sex) | 240.300672 | 1 | 2.512809 | 0.125494 |
| max_path_length:Age | 278.652103 | 1 | 2.913847 | 0.100215 |
| max_path_length:Age:C(Sex) | 353.063345 | 1 | 3.691961 | 0.066146 |
| Residual | 2390.757787 | 25 | NaN | NaN |

**d.** CFPT UP

| factors | sum_sq | df | F | PR(>F) |
|---|---|---|---|---|
| C(Sex) | 9.362467 | 1 | 0.04161 | 0.840017 |
| max_path_length | 34.833795 | 1 | 0.154812 | 0.697314 |
| max_path_length:C(Sex) | 206.915664 | 1 | 0.919598 | 0.346764 |
| Age | 2.292185 | 1 | 0.010187 | 0.92041 |
| Age:C(Sex) | 453.880903 | 1 | 2.017189 | 0.167879 |
| max_path_length:Age | 65.350725 | 1 | 0.290439 | 0.594706 |
| max_path_length:Age:C(Sex) | 2.09508 | 1 | 0.009311 | 0.923897 |
| Residual | 5625.165614 | 25 | NaN | NaN |

**c.** CHMT

| factors | sum_sq | df | F | PR(>F) |
|---|---|---|---|---|
| C(Sex) | 31.150909 | 1 | 1.755536 | 0.197166 |
| max_path_length | 17.086768 | 1 | 0.962939 | 0.335849 |
| max_path_length:C(Sex) | 26.486223 | 1 | 1.492654 | 0.233201 |
| Age | 0.056269 | 1 | 0.003171 | 0.95554 |
| Age:C(Sex) | 47.246254 | 1 | 2.662603 | 0.115267 |
| max_path_length:Age | 19.186108 | 1 | 1.08125 | 0.308371 |
| max_path_length:Age:C(Sex) | 4.307978 | 1 | 0.24278 | 0.626505 |
| Residual | 443.609596 | 25 | NaN | NaN |

**d.** Eyes Test emotion

| factors | sum_sq | df | F | PR(>F) |
|---|---|---|---|---|
| C(Sex) | 32.865961 | 1 | 10.265191 | 0.00368 |
| max_path_length | 5.490531 | 1 | 1.714885 | 0.202261 |
| max_path_length:C(Sex) | 0.66925 | 1 | 0.20903 | 0.651477 |
| Age | 3.484016 | 1 | 1.08818 | 0.306858 |
| Age:C(Sex) | 73.186508 | 1 | 22.858711 | 0.000066 |
| max_path_length:Age | 20.306577 | 1 | 6.342456 | 0.018558 |
| max_path_length:Age:C(Sex) | 32.404479 | 1 | 10.121054 | 0.003889 |
| Residual | 80.04225 | 25 | NaN | NaN |

**e.** Famous Faces UK Exposed

| factors | sum_sq | df | F | PR(>F) |
|---|---|---|---|---|
| C(Sex) | 0.277592 | 1 | 0.023709 | 0.878864 |
| max_path_length | 0.002829 | 1 | 0.000242 | 0.987721 |
| max_path_length:C(Sex) | 2.928159 | 1 | 0.250091 | 0.621385 |
| Age | 0.863326 | 1 | 0.073736 | 0.788203 |
| Age:C(Sex) | 0.080393 | 1 | 0.006866 | 0.93462 |
| max_path_length:Age | 19.138601 | 1 | 1.634605 | 0.21281 |
| max_path_length:Age:C(Sex) | 18.97919 | 1 | 1.62099 | 0.214666 |
| Residual | 292.709919 | 25 | NaN | NaN |

**f.** Famous Faces UK Identified

| factors | sum_sq | df | F | PR(>F) |
|---|---|---|---|---|
| C(Sex) | 47.097795 | 1 | 0.840144 | 0.368113 |
| max_path_length | 46.545723 | 1 | 0.830296 | 0.37089 |
| max_path_length:C(Sex) | 53.481449 | 1 | 0.954017 | 0.338056 |
| Age | 100.18095 | 1 | 1.787056 | 0.193324 |
| Age:C(Sex) | 83.860182 | 1 | 1.495922 | 0.232704 |
| max_path_length:Age | 86.914648 | 1 | 1.550408 | 0.22462 |
| max_path_length:Age:C(Sex) | 82.907967 | 1 | 1.478936 | 0.235299 |
| Residual | 1401.479943 | 25 | NaN | NaN |

**g.** Films Task Up

| factors | sum_sq | df | F | PR(>F) |
|---|---|---|---|---|
| C(Sex) | 232.202904 | 1 | 7.240522 | 0.012521 |
| max_path_length | 14.767653 | 1 | 0.460483 | 0.503632 |
| max_path_length:C(Sex) | 60.761522 | 1 | 1.894658 | 0.180882 |
| Age | 7.189642 | 1 | 0.224187 | 0.639977 |
| Age:C(Sex) | 205.047001 | 1 | 6.39375 | 0.018138 |
| max_path_length:Age | 16.17416 | 1 | 0.504341 | 0.484171 |
| max_path_length:Age:C(Sex) | 74.323123 | 1 | 2.317534 | 0.140472 |
| Residual | 801.747808 | 25 | NaN | NaN |

**h.** WASI Full

| factors | sum_sq | df | F | PR(>F) |
|---|---|---|---|---|
| C(Sex) | 47.718514 | 1 | 1.112217 | 0.303584 |
| max_path_length | 26.802479 | 1 | 0.624709 | 0.438138 |
| max_path_length:C(Sex) | 91.549122 | 1 | 2.133815 | 0.158883 |
| Age | 58.781486 | 1 | 1.370071 | 0.254912 |
| Age:C(Sex) | 31.348575 | 1 | 0.730668 | 0.402314 |
| max_path_length:Age | 35.797524 | 1 | 0.834364 | 0.371382 |
| max_path_length:Age:C(Sex) | 78.862339 | 1 | 1.838113 | 0.189574 |
| Residual | 900.983407 | 21 | NaN | NaN |

**i.** WASI Verbal

| factors | sum_sq | df | F | PR(>F) |
|---|---|---|---|---|
| C(Sex) | 18.655828 | 1 | 0.220149 | 0.643762 |
| max_path_length | 73.829691 | 1 | 0.871229 | 0.361224 |
| max_path_length:C(Sex) | 72.560332 | 1 | 0.85625 | 0.365303 |
| Age | 11.006257 | 1 | 0.12988 | 0.722156 |
| Age:C(Sex) | 22.838643 | 1 | 0.269508 | 0.60909 |
| max_path_length:Age | 0.509091 | 1 | 0.006008 | 0.938953 |
| max_path_length:Age:C(Sex) | 17.410809 | 1 | 0.205457 | 0.655001 |
| Residual | 1779.58207 | 21 | NaN | NaN |

**j.** WASI Performance

| factors | sum_sq | df | F | PR(>F) |
|---|---|---|---|---|
| C(Sex) | 40.166612 | 1 | 0.811282 | 0.377955 |
| max_path_length | 0.080598 | 1 | 0.001628 | 0.968197 |
| max_path_length:C(Sex) | 70.294943 | 1 | 1.419811 | 0.246728 |
| Age | 99.888311 | 1 | 2.017536 | 0.170169 |
| Age:C(Sex) | 13.268875 | 1 | 0.268004 | 0.610085 |
| max_path_length:Age | 92.885624 | 1 | 1.876096 | 0.185248 |
| max_path_length:Age:C(Sex) | 134.342902 | 1 | 2.713447 | 0.114389 |
| Residual | 1039.711221 | 21 | NaN | NaN |

**a.** Horses

| factors | sum_sq | df | F | PR(>F) |
|---|---|---|---|---|
| C(Sex) | 0.00069 | 1 | 0.225546 | 0.638969 |
| max_path_length | 0.002536 | 1 | 0.829437 | 0.371134 |
| max_path_length:C(Sex) | 0.000142 | 1 | 0.046537 | 0.830953 |
| Age | 0.000621 | 1 | 0.203199 | 0.656037 |
| Age:C(Sex) | 0.001296 | 1 | 0.42392 | 0.520928 |
| max_path_length:Age | 0.002146 | 1 | 0.701669 | 0.41016 |
| max_path_length:Age:C(Sex) | 0.001585 | 1 | 0.518483 | 0.478168 |
| Residual | 0.076447 | 25 | NaN | NaN |

**b.** Horses RT

| factors | sum_sq | df | F | PR(>F) |
|---|---|---|---|---|
| C(Sex) | 999429.78758 | 1 | 12.385358 | 0.001682 |
| max_path_length | 26095.263241 | 1 | 0.323384 | 0.574655 |
| max_path_length:C(Sex) | 261554.415422 | 1 | 3.241293 | 0.083883 |
| Age | 261966.484605 | 1 | 3.2464 | 0.083653 |
| Age:C(Sex) | 410008.53263 | 1 | 5.081 | 0.033203 |
| max_path_length:Age | 48299.132328 | 1 | 0.598543 | 0.446389 |
| max_path_length:Age:C(Sex) | 17903.249388 | 1 | 0.221865 | 0.641707 |
| Residual | 2017361.60179 | 25 | NaN | NaN |

**c.** Sequential Em Inv

| factors | sum_sq | df | F | PR(>F) |
|---|---|---|---|---|
| C(Sex) | 41.954007 | 1 | 1.660215 | 0.209372 |
| max_path_length | 2.770039 | 1 | 0.109617 | 0.74334 |
| max_path_length:C(Sex) | 10.295551 | 1 | 0.407418 | 0.529087 |
| Age | 11.892025 | 1 | 0.470594 | 0.499027 |
| Age:C(Sex) | 50.089906 | 1 | 1.982171 | 0.171476 |
| max_path_length:Age | 1.723653 | 1 | 0.068209 | 0.796102 |
| max_path_length:Age:C(Sex) | 0.133138 | 1 | 0.005269 | 0.942714 |
| Residual | 631.755664 | 25 | NaN | NaN |

**d.** Sequential Em emup

| factors | sum_sq | df | F | PR(>F) |
|---|---|---|---|---|
| C(Sex) | 7.146251 | 1 | 0.447116 | 0.509836 |
| max_path_length | 46.44166 | 1 | 2.905691 | 0.100666 |
| max_path_length:C(Sex) | 103.879453 | 1 | 6.49937 | 0.017305 |
| Age | 67.012933 | 1 | 4.192762 | 0.051241 |
| Age:C(Sex) | 90.620263 | 1 | 5.66979 | 0.025192 |
| max_path_length:Age | 28.259287 | 1 | 1.768084 | 0.195625 |
| max_path_length:Age:C(Sex) | 64.714013 | 1 | 4.048927 | 0.055092 |
| Residual | 399.575067 | 25 | NaN | NaN |

**c.** Sequential ID Inv

| factors | sum_sq | df | F | PR(>F) |
|---|---|---|---|---|
| C(Sex) | 0.422975 | 1 | 0.02783 | 0.868852 |
| max_path_length | 26.245187 | 1 | 1.726807 | 0.20075 |
| max_path_length:C(Sex) | 0.116773 | 1 | 0.007683 | 0.93085 |
| Age | 173.480469 | 1 | 11.414181 | 0.002391 |
| Age:C(Sex) | 0.148093 | 1 | 0.009744 | 0.922155 |
| max_path_length:Age | 0.650911 | 1 | 0.042827 | 0.837728 |
| max_path_length:Age:C(Sex) | 0.74763 | 1 | 0.04919 | 0.82628 |
| Residual | 379.966973 | 25 | NaN | NaN |

**d.** Sequential ID up

| factors | sum_sq | df | F | PR(>F) |
|---|---|---|---|---|
| C(Sex) | 19.029644 | 1 | 1.30097 | 0.264846 |
| max_path_length | 19.510609 | 1 | 1.333851 | 0.259041 |
| max_path_length:C(Sex) | 12.58389 | 1 | 0.860303 | 0.362523 |
| Age | 79.607421 | 1 | 5.442395 | 0.027999 |
| Age:C(Sex) | 1.019314 | 1 | 0.069686 | 0.793959 |
| max_path_length:Age | 0.210603 | 1 | 0.014398 | 0.905449 |
| max_path_length:Age:C(Sex) | 21.772067 | 1 | 1.488456 | 0.23384 |
| Residual | 365.681946 | 25 | NaN | NaN |

**e.** PC1

| factors | sum_sq | df | F | PR(>F) |
|---|---|---|---|---|
| C(Sex) | 0.111932 | 1 | 0.260882 | 0.613996 |
| max_path_length | 1.449289 | 1 | 3.377896 | 0.077989 |
| max_path_length:C(Sex) | 0.184643 | 1 | 0.430352 | 0.517809 |
| Age | 0.065715 | 1 | 0.153163 | 0.698846 |
| Age:C(Sex) | 0.296455 | 1 | 0.690955 | 0.413709 |
| max_path_length:Age | 0.101779 | 1 | 0.237218 | 0.630466 |
| max_path_length:Age:C(Sex) | 0.86811 | 1 | 2.023328 | 0.167258 |
| Residual | 10.72627 | 25 | NaN | NaN |

**f.** PC2

| factors | sum_sq | df | F | PR(>F) |
|---|---|---|---|---|
| C(Sex) | 0.684603 | 1 | 0.580594 | 0.453209 |
| max_path_length | 2.362286 | 1 | 2.003394 | 0.169285 |
| max_path_length:C(Sex) | 1.944911 | 1 | 1.649429 | 0.210811 |
| Age | 0.120878 | 1 | 0.102513 | 0.751495 |
| Age:C(Sex) | 0.028433 | 1 | 0.024113 | 0.877844 |
| max_path_length:Age | 0.029028 | 1 | 0.024618 | 0.876583 |
| max_path_length:Age:C(Sex) | 1.728593 | 1 | 1.465975 | 0.237304 |
| Residual | 29.47856 | 25 | NaN | NaN |

**g.** PC3

| factors | sum_sq | df | F | PR(>F) |
|---|---|---|---|---|
| C(Sex) | 7.375231 | 1 | 11.815077 | 0.002065 |
| max_path_length | 0.08111 | 1 | 0.129938 | 0.721525 |
| max_path_length:C(Sex) | 1.444492 | 1 | 2.314067 | 0.140757 |
| Age | 0.115526 | 1 | 0.185072 | 0.670736 |
| Age:C(Sex) | 11.868392 | 1 | 19.013093 | 0.000196 |
| max_path_length:Age | 0.003847 | 1 | 0.006163 | 0.938054 |
| max_path_length:Age:C(Sex) | 1.199902 | 1 | 1.922236 | 0.177852 |
| Residual | 15.605551 | 25 | NaN | NaN |

**h.** Cars RT

| factors | sum_sq | df | F | PR(>F) |
|---|---|---|---|---|
| C(Sex) | 154598.055858 | 1 | 1.161978 | 0.291347 |
| max_path_length | 4310.694698 | 1 | 0.0324 | 0.858604 |
| max_path_length:C(Sex) | 20415.707205 | 1 | 0.153447 | 0.698582 |
| Age | 490507.131405 | 1 | 3.686712 | 0.066327 |
| Age:C(Sex) | 494604.907649 | 1 | 3.717511 | 0.065277 |
| max_path_length:Age | 519421.079619 | 1 | 3.904032 | 0.059306 |
| max_path_length:Age:C(Sex) | 242121.510236 | 1 | 1.819815 | 0.189428 |
| Residual | 3326183.136101 | 25 | NaN | NaN |

**i.** Cars

| factors | sum_sq | df | F | PR(>F) |
|---|---|---|---|---|
| C(Sex) | 0.025823 | 1 | 5.44236 | 0.027999 |
| max_path_length | 0.004492 | 1 | 0.946697 | 0.339882 |
| max_path_length:C(Sex) | 0.003729 | 1 | 0.785967 | 0.383771 |
| Age | 0.005242 | 1 | 1.10479 | 0.303272 |
| Age:C(Sex) | 0.001312 | 1 | 0.276417 | 0.603692 |
| max_path_length:Age | 0.000044 | 1 | 0.009312 | 0.923896 |
| max_path_length:Age:C(Sex) | 0.008402 | 1 | 1.770705 | 0.195305 |
| Residual | 0.11862 | 25 | NaN | NaN |

**j.**

**Supplementary Figure 2.2 Testing for a relationship between MFS length and other behavioral tests in Dataset 1 in DPs: No significant effects.** Same layout as Supplementary Figure 2.1, but for DPs only. The relationship between MFS length and behavior was specific to CFMT performance.

**a.** Baldwomen

| factors | sum_sq | df | F | PR(>F) |
|---|---|---|---|---|
| C(Sex) | 0.000266 | 1 | 1.049303 | 0.314437 |
| max_path_length | 0.000048 | 1 | 0.190717 | 0.66567 |
| max_path_length:C(Sex) | 0.000063 | 1 | 0.249142 | 0.621579 |
| Age | 0.000124 | 1 | 0.489 | 0.490143 |
| Age:C(Sex) | 0.001741 | 1 | 6.869889 | 0.014009 |
| max_path_length:Age | 0.000057 | 1 | 0.225992 | 0.638199 |
| max_path_length:Age:C(Sex) | 0.000129 | 1 | 0.507167 | 0.48226 |
| Residual | 0.007098 | 28 | NaN | NaN |

**b.** Baldwomen RT

| factors | sum_sq | df | F | PR(>F) |
|---|---|---|---|---|
| C(Sex) | 754146.835922 | 1 | 30.213835 | 0.000007 |
| max_path_length | 3386.311642 | 1 | 0.135668 | 0.715397 |
| max_path_length:C(Sex) | 37619.22763 | 1 | 1.507162 | 0.229794 |
| Age | 826.642841 | 1 | 0.033118 | 0.856906 |
| Age:C(Sex) | 235207.26551 | 1 | 9.423249 | 0.004724 |
| max_path_length:Age | 3035.74755 | 1 | 0.121623 | 0.72989 |
| max_path_length:Age:C(Sex) | 1742.500251 | 1 | 0.069811 | 0.793548 |
| Residual | 698888.805705 | 28 | NaN | NaN |

**c.** CFPT INV

| factors | sum_sq | df | F | PR(>F) |
|---|---|---|---|---|
| C(Sex) | 768.493918 | 1 | 2.842824 | 0.102896 |
| max_path_length | 288.841162 | 1 | 1.068485 | 0.310134 |
| max_path_length:C(Sex) | 296.445831 | 1 | 1.096617 | 0.303966 |
| Age | 1.941889 | 1 | 0.007183 | 0.933059 |
| Age:C(Sex) | 1444.669379 | 1 | 5.344142 | 0.028366 |
| max_path_length:Age | 237.46259 | 1 | 0.878425 | 0.356648 |
| max_path_length:Age:C(Sex) | 69.781295 | 1 | 0.258136 | 0.615383 |
| Residual | 7569.174942 | 28 | NaN | NaN |

**d.** CFPT UP

| factors | sum_sq | df | F | PR(>F) |
|---|---|---|---|---|
| C(Sex) | 165.110141 | 1 | 1.176555 | 0.28731 |
| max_path_length | 713.956214 | 1 | 5.087566 | 0.0321 |
| max_path_length:C(Sex) | 24.716033 | 1 | 0.176123 | 0.677929 |
| Age | 187.942128 | 1 | 1.339253 | 0.256943 |
| Age:C(Sex) | 29.796971 | 1 | 0.21233 | 0.648507 |
| max_path_length:Age | 2.308727 | 1 | 0.016452 | 0.898857 |
| max_path_length:Age:C(Sex) | 24.116972 | 1 | 0.171855 | 0.681628 |
| Residual | 3929.339512 | 28 | NaN | NaN |

**c.** CHMT

| factors | sum_sq | df | F | PR(>F) |
|---|---|---|---|---|
| C(Sex) | 43.010474 | 1 | 2.212594 | 0.148064 |
| max_path_length | 4.748272 | 1 | 0.244266 | 0.624997 |
| max_path_length:C(Sex) | 25.085807 | 1 | 1.290493 | 0.265587 |
| Age | 2.070542 | 1 | 0.106515 | 0.746574 |
| Age:C(Sex) | 10.478337 | 1 | 0.539039 | 0.468936 |
| max_path_length:Age | 2.939948 | 1 | 0.15124 | 0.700295 |
| max_path_length:Age:C(Sex) | 73.794908 | 1 | 3.796242 | 0.061454 |
| Residual | 544.290159 | 28 | NaN | NaN |

**d.** Eyes Test emotion

| factors | sum_sq | df | F | PR(>F) |
|---|---|---|---|---|
| C(Sex) | 14.88746 | 1 | 2.030001 | 0.165272 |
| max_path_length | 2.711648 | 1 | 0.369751 | 0.548041 |
| max_path_length:C(Sex) | 0.124551 | 1 | 0.016983 | 0.897245 |
| Age | 13.845981 | 1 | 1.887989 | 0.180329 |
| Age:C(Sex) | 13.357249 | 1 | 1.821347 | 0.187964 |
| max_path_length:Age | 0.353177 | 1 | 0.048158 | 0.827892 |
| max_path_length:Age:C(Sex) | 0.001805 | 1 | 0.000246 | 0.987594 |
| Residual | 205.344189 | 28 | NaN | NaN |

**e.** Famous Faces UK Exposed

| factors | sum_sq | df | F | PR(>F) |
|---|---|---|---|---|
| C(Sex) | 5.592898 | 1 | 0.153624 | 0.698063 |
| max_path_length | 12.57756 | 1 | 0.345477 | 0.561395 |
| max_path_length:C(Sex) | 67.072311 | 1 | 1.842326 | 0.185519 |
| Age | 27.86325 | 1 | 0.765341 | 0.389105 |
| Age:C(Sex) | 1.026957 | 1 | 0.028208 | 0.867827 |
| max_path_length:Age | 6.65426 | 1 | 0.182778 | 0.672267 |
| max_path_length:Age:C(Sex) | 10.11595 | 1 | 0.277863 | 0.602258 |
| Residual | 1019.376742 | 28 | NaN | NaN |

**f.** Famous Faces UK Identified

| factors | sum_sq | df | F | PR(>F) |
|---|---|---|---|---|
| C(Sex) | 12.820373 | 1 | 0.192982 | 0.663818 |
| max_path_length | 4.970403 | 1 | 0.074818 | 0.786454 |
| max_path_length:C(Sex) | 296.250516 | 1 | 4.459396 | 0.043763 |
| Age | 81.631004 | 1 | 1.228774 | 0.277077 |
| Age:C(Sex) | 113.366954 | 1 | 1.706489 | 0.202071 |
| max_path_length:Age | 47.832554 | 1 | 0.720013 | 0.403336 |
| max_path_length:Age:C(Sex) | 92.548649 | 1 | 1.393115 | 0.24781 |
| Residual | 1860.120529 | 28 | NaN | NaN |

**g.** Films Task Up

| factors | sum_sq | df | F | PR(>F) |
|---|---|---|---|---|
| C(Sex) | 2.740356 | 1 | 0.078073 | 0.78198 |
| max_path_length | 29.478326 | 1 | 0.839841 | 0.367268 |
| max_path_length:C(Sex) | 0.639328 | 1 | 0.018215 | 0.893608 |
| Age | 63.824324 | 1 | 1.818363 | 0.188315 |
| Age:C(Sex) | 24.178675 | 1 | 0.688853 | 0.413575 |
| max_path_length:Age | 0.170323 | 1 | 0.004853 | 0.944959 |
| max_path_length:Age:C(Sex) | 20.166156 | 1 | 0.574536 | 0.454795 |
| Residual | 982.796721 | 28 | NaN | NaN |

**h.** WASI Full

| factors | sum_sq | df | F | PR(>F) |
|---|---|---|---|---|
| C(Sex) | 16.50774 | 1 | 0.286538 | 0.596674 |
| max_path_length | 165.0984 | 1 | 2.865743 | 0.101584 |
| max_path_length:C(Sex) | 0.28581 | 1 | 0.004961 | 0.944348 |
| Age | 33.804423 | 1 | 0.58677 | 0.450081 |
| Age:C(Sex) | 487.786606 | 1 | 8.466896 | 0.007013 |
| max_path_length:Age | 14.295546 | 1 | 0.248139 | 0.622279 |
| max_path_length:Age:C(Sex) | 169.686732 | 1 | 2.945386 | 0.097169 |
| Residual | 1613.108788 | 28 | NaN | NaN |

**i.** WASI Verbal

| factors | sum_sq | df | F | PR(>F) |
|---|---|---|---|---|
| C(Sex) | 16.857332 | 1 | 0.121915 | 0.729579 |
| max_path_length | 205.345588 | 1 | 1.485094 | 0.233148 |
| max_path_length:C(Sex) | 1.663428 | 1 | 0.01203 | 0.913444 |
| Age | 77.789048 | 1 | 0.562583 | 0.459478 |
| Age:C(Sex) | 586.374584 | 1 | 4.240759 | 0.048873 |
| max_path_length:Age | 124.022406 | 1 | 0.896951 | 0.351703 |
| max_path_length:Age:C(Sex) | 286.459352 | 1 | 2.071722 | 0.161139 |
| Residual | 3871.592125 | 28 | NaN | NaN |

**j.** WASI Performance

| factors | sum_sq | df | F | PR(>F) |
|---|---|---|---|---|
| C(Sex) | 114.188435 | 1 | 3.322481 | 0.079034 |
| max_path_length | 53.969865 | 1 | 1.570333 | 0.220523 |
| max_path_length:C(Sex) | 1.387067 | 1 | 0.040359 | 0.842233 |
| Age | 8.829761 | 1 | 0.256915 | 0.616216 |
| Age:C(Sex) | 182.778252 | 1 | 5.318204 | 0.028721 |
| max_path_length:Age | 24.001084 | 1 | 0.698347 | 0.410414 |
| max_path_length:Age:C(Sex) | 15.581778 | 1 | 0.453375 | 0.506257 |
| Residual | 962.315744 | 28 | NaN | NaN |

**a.** Horses

| factors | sum_sq | df | F | PR(>F) |
|---|---|---|---|---|
| C(Sex) | 0.00112 | 1 | 0.67437 | 0.418469 |
| max_path_length | 0 | 1 | 0.000144 | 0.990523 |
| max_path_length:C(Sex) | 0.00035 | 1 | 0.210835 | 0.649659 |
| Age | 0.000529 | 1 | 0.318668 | 0.576904 |
| Age:C(Sex) | 0.012588 | 1 | 7.578676 | 0.01025 |
| max_path_length:Age | 0.000051 | 1 | 0.030857 | 0.861824 |
| max_path_length:Age:C(Sex) | 0.000075 | 1 | 0.045193 | 0.83319 |
| Residual | 0.046506 | 28 | NaN | NaN |

**b.** Horses RT

| factors | sum_sq | df | F | PR(>F) |
|---|---|---|---|---|
| C(Sex) | 950162.067779 | 1 | 16.821846 | 0.00032 |
| max_path_length | 28920.783294 | 1 | 0.512019 | 0.480191 |
| max_path_length:C(Sex) | 84911.432065 | 1 | 1.503288 | 0.230378 |
| Age | 204.938726 | 1 | 0.003628 | 0.952396 |
| Age:C(Sex) | 1225474.573363 | 1 | 21.696029 | 0.000071 |
| max_path_length:Age | 10334.50875 | 1 | 0.182964 | 0.67211 |
| max_path_length:Age:C(Sex) | 6943.490974 | 1 | 0.122929 | 0.728504 |
| Residual | 1581546.909796 | 28 | NaN | NaN |

**c.** Sequential Em Inv

| factors | sum_sq | df | F | PR(>F) |
|---|---|---|---|---|
| C(Sex) | 96.912855 | 1 | 2.121329 | 0.156384 |
| max_path_length | 22.819603 | 1 | 0.499499 | 0.485561 |
| max_path_length:C(Sex) | 6.363355 | 1 | 0.139288 | 0.711801 |
| Age | 1.369656 | 1 | 0.02998 | 0.86378 |
| Age:C(Sex) | 19.926743 | 1 | 0.436177 | 0.51437 |
| max_path_length:Age | 23.699064 | 1 | 0.51875 | 0.477346 |
| max_path_length:Age:C(Sex) | 17.822367 | 1 | 0.390114 | 0.537296 |
| Residual | 1279.179321 | 28 | NaN | NaN |

**d.** Sequential Em emup

| factors | sum_sq | df | F | PR(>F) |
|---|---|---|---|---|
| C(Sex) | 0.211676 | 1 | 0.014633 | 0.904581 |
| max_path_length | 12.634015 | 1 | 0.873373 | 0.358013 |
| max_path_length:C(Sex) | 12.780658 | 1 | 0.88351 | 0.355281 |
| Age | 32.332506 | 1 | 2.235104 | 0.146094 |
| Age:C(Sex) | 0.707601 | 1 | 0.048916 | 0.826566 |
| max_path_length:Age | 3.31613 | 1 | 0.22924 | 0.635806 |
| max_path_length:Age:C(Sex) | 2.750127 | 1 | 0.190113 | 0.666166 |
| Residual | 405.041623 | 28 | NaN | NaN |

**c.** Sequential ID Inv

| factors | sum_sq | df | F | PR(>F) |
|---|---|---|---|---|
| C(Sex) | 1.150229 | 1 | 0.080759 | 0.778363 |
| max_path_length | 6.900187 | 1 | 0.484471 | 0.492142 |
| max_path_length:C(Sex) | 14.054085 | 1 | 0.986755 | 0.329044 |
| Age | 240.643228 | 1 | 16.895859 | 0.000312 |
| Age:C(Sex) | 4.387346 | 1 | 0.308041 | 0.583292 |
| max_path_length:Age | 0.132247 | 1 | 0.009285 | 0.923921 |
| max_path_length:Age:C(Sex) | 1.28946 | 1 | 0.090535 | 0.765722 |
| Residual | 398.796546 | 28 | NaN | NaN |

**d.** Sequential ID up

| factors | sum_sq | df | F | PR(>F) |
|---|---|---|---|---|
| C(Sex) | 86.76178 | 1 | 2.700086 | 0.111527 |
| max_path_length | 84.647415 | 1 | 2.634285 | 0.115786 |
| max_path_length:C(Sex) | 10.541286 | 1 | 0.328052 | 0.57138 |
| Age | 50.385262 | 1 | 1.568024 | 0.220853 |
| Age:C(Sex) | 165.246104 | 1 | 5.142572 | 0.031256 |
| max_path_length:Age | 0.027593 | 1 | 0.000859 | 0.97683 |
| max_path_length:Age:C(Sex) | 3.859116 | 1 | 0.120098 | 0.731519 |
| Residual | 899.723178 | 28 | NaN | NaN |

**e.** PC1

| factors | sum_sq | df | F | PR(>F) |
|---|---|---|---|---|
| C(Sex) | 0.137934 | 1 | 0.645931 | 0.428342 |
| max_path_length | 0.294004 | 1 | 1.376787 | 0.250534 |
| max_path_length:C(Sex) | 0.004888 | 1 | 0.02289 | 0.880828 |
| Age | 0.249675 | 1 | 1.169201 | 0.288791 |
| Age:C(Sex) | 0.037141 | 1 | 0.173928 | 0.679824 |
| max_path_length:Age | 0.075902 | 1 | 0.355439 | 0.55584 |
| max_path_length:Age:C(Sex) | 0.068779 | 1 | 0.322086 | 0.574879 |
| Residual | 5.979223 | 28 | NaN | NaN |

**f.** PC2

| factors | sum_sq | df | F | PR(>F) |
|---|---|---|---|---|
| C(Sex) | 0.688417 | 1 | 2.281506 | 0.142129 |
| max_path_length | 0.211489 | 1 | 0.700903 | 0.40957 |
| max_path_length:C(Sex) | 0.004715 | 1 | 0.015627 | 0.90141 |
| Age | 0.322354 | 1 | 1.068325 | 0.310169 |
| Age:C(Sex) | 0.543957 | 1 | 1.802746 | 0.190165 |
| max_path_length:Age | 0.002552 | 1 | 0.008456 | 0.927387 |
| max_path_length:Age:C(Sex) | 0.214035 | 1 | 0.709339 | 0.4068 |
| Residual | 8.448667 | 28 | NaN | NaN |

**g.** PC3

| factors | sum_sq | df | F | PR(>F) |
|---|---|---|---|---|
| C(Sex) | 0.142809 | 1 | 0.177789 | 0.6765 |
| max_path_length | 0.063018 | 1 | 0.078454 | 0.781462 |
| max_path_length:C(Sex) | 0.010851 | 1 | 0.013509 | 0.908301 |
| Age | 0.031889 | 1 | 0.039701 | 0.843507 |
| Age:C(Sex) | 0.884118 | 1 | 1.100679 | 0.303089 |
| max_path_length:Age | 0.004102 | 1 | 0.005107 | 0.943538 |
| max_path_length:Age:C(Sex) | 0.109552 | 1 | 0.136386 | 0.714679 |
| Residual | 22.490933 | 28 | NaN | NaN |

**h.** Cars RT

| factors | sum_sq | df | F | PR(>F) |
|---|---|---|---|---|
| C(Sex) | 44249.190663 | 1 | 1.858196 | 0.183695 |
| max_path_length | 865.042008 | 1 | 0.036326 | 0.850218 |
| max_path_length:C(Sex) | 10857.760666 | 1 | 0.45596 | 0.505057 |
| Age | 42457.983157 | 1 | 1.782976 | 0.192539 |
| Age:C(Sex) | 64018.448972 | 1 | 2.688384 | 0.112271 |
| max_path_length:Age | 16.991162 | 1 | 0.000714 | 0.978879 |
| max_path_length:Age:C(Sex) | 10010.02284 | 1 | 0.42036 | 0.52204 |
| Residual | 666763.63274 | 28 | NaN | NaN |

**i.** Cars

| factors | sum_sq | df | F | PR(>F) |
|---|---|---|---|---|
| C(Sex) | 0.000041 | 1 | 0.044353 | 0.834724 |
| max_path_length | 0.000073 | 1 | 0.07996 | 0.779431 |
| max_path_length:C(Sex) | 0.000415 | 1 | 0.45186 | 0.506963 |
| Age | 0.015993 | 1 | 17.409756 | 0.000264 |
| Age:C(Sex) | 0.00061 | 1 | 0.664381 | 0.421897 |
| max_path_length:Age | 0.000304 | 1 | 0.331021 | 0.569654 |
| max_path_length:Age:C(Sex) | 0.000111 | 1 | 0.120859 | 0.730705 |
| Residual | 0.025721 | 28 | NaN | NaN |

**j.**

**Supplementary Figure 2.3 Testing for a relationship between MFS length and other behavioral tests in Dataset 1 in NTs: No significant effects.** Same layout as Supplementary Figure 2.1, but for NTs only. The relationship between MFS length and behavior was specific to CFMT performance.

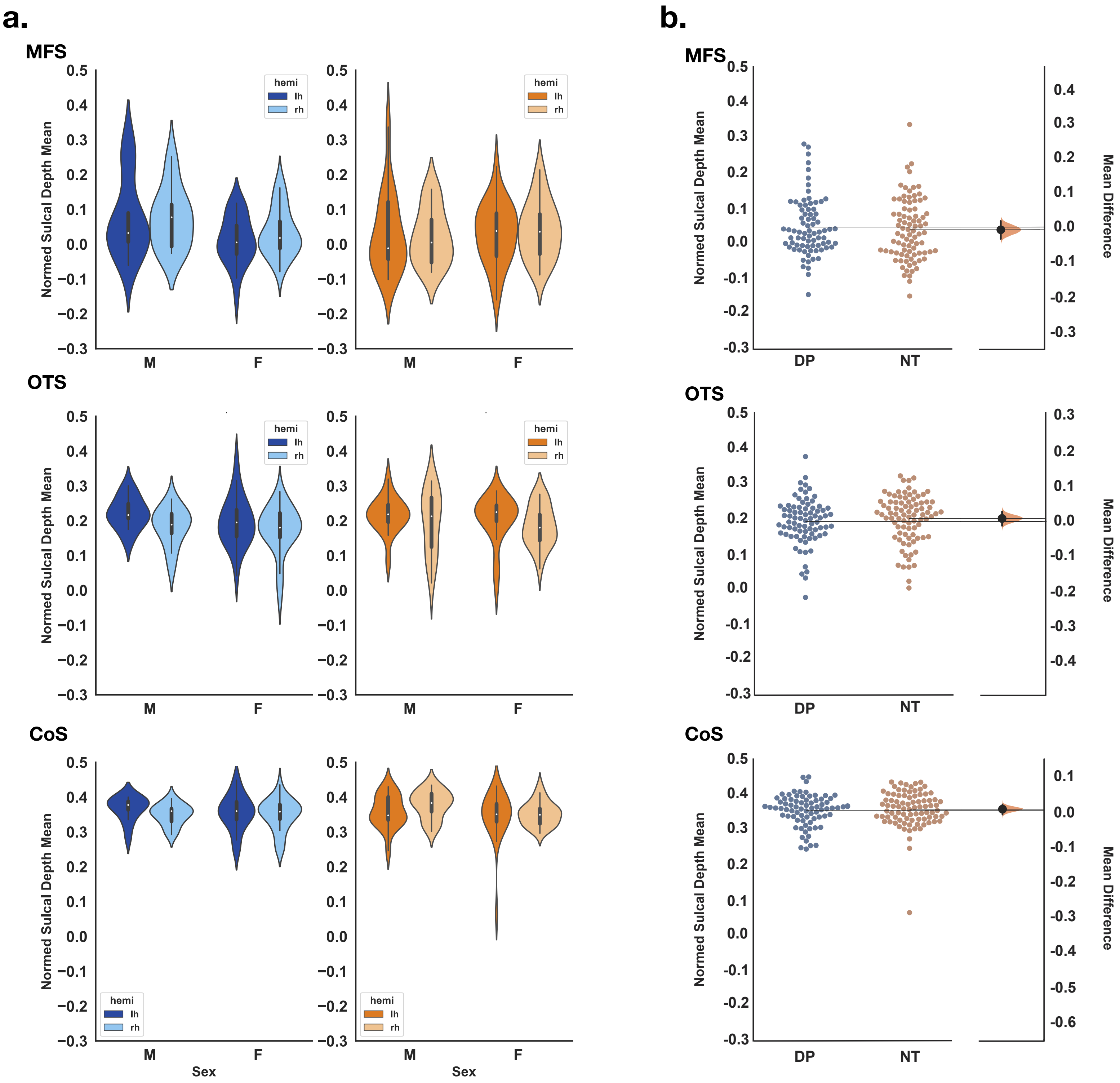

**Supplementary Figure 3. Depth of the MFS, OTS, and CoS do not differ between DPs and NTs. a.** Violin plots depicting the mean sulcal depth normalized to the maximum depth across the cerebral cortex (DPs, blue; NTs, orange; lh, darker shade; rh, lighter shade). As expected, the MFS was the shallowest sulcus, with an average normed mean depth of 0.03±0.09. **b.** Swarm plot showing sulcal depth as a function of group (DPs, blue; NTs, orange). On the right y-axis, a bootstrap of 5,000 iterations was used to generate a confidence interval (black bar) displayed within density plot (orange) which depicts the mean difference in sulcal depth between groups for each iteration. None of the sulci significantly differed between DPs and NTs ($F(1,468) < .001$, $p > .97$).

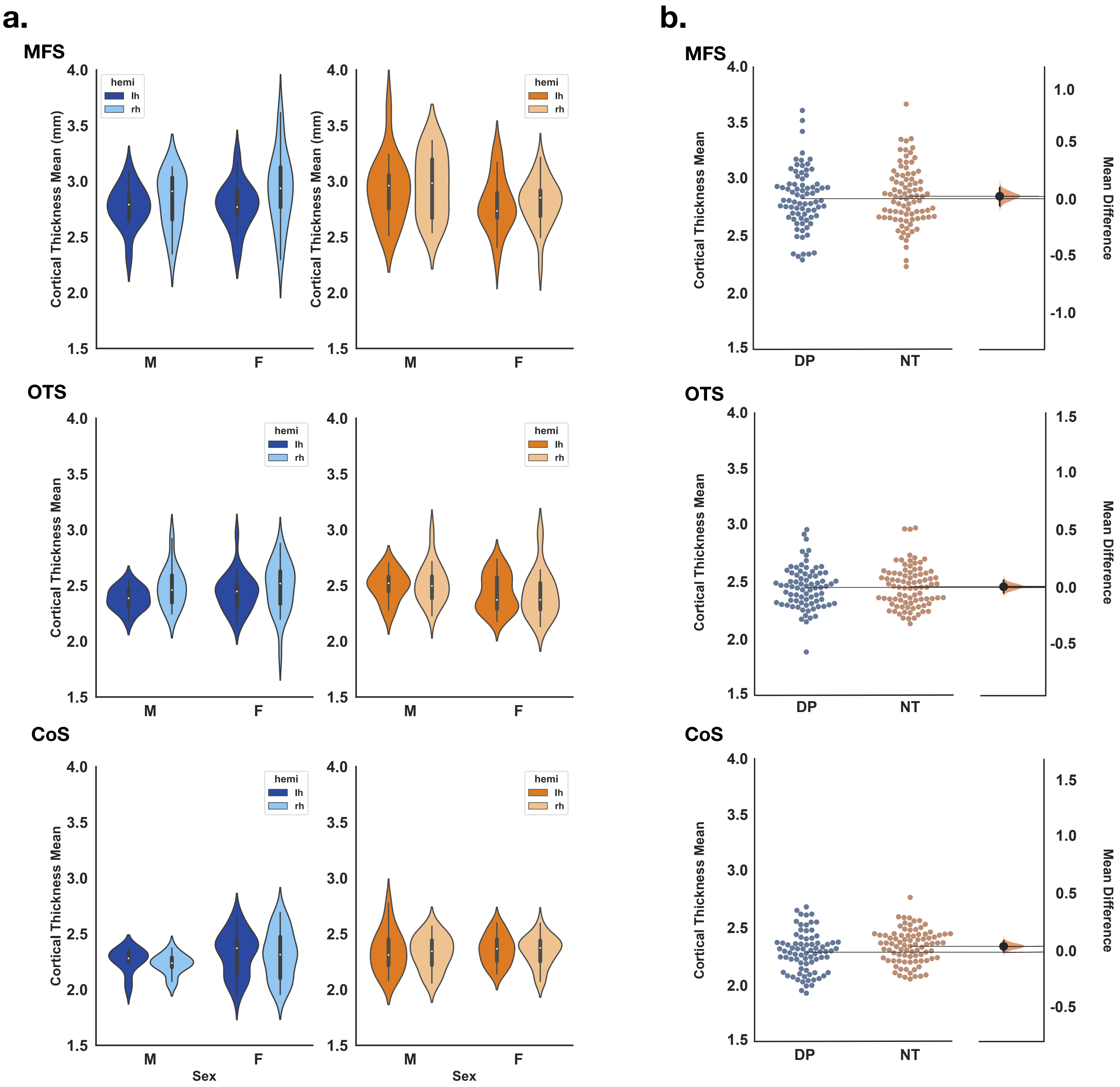

**Supplementary Figure 4. Cortical thickness of the MFS, OTS, and CoS does not differ between DPs and NTs. a.** Violin plots depicting the cortical thickness (CT) by group (DPs, blue; NTs, orange), hemisphere (lh, darker shade; rh, lighter shade), and sulcus. Of the three VTC sulci, the MFS had the greatest CT (2.84±0.26mm). **b.** Swarm plot depicting CT as a function of group (DPs, blue; NTs, orange). On the right y-axis, a bootstrap of 5,000 iterations was used to generate a confidence interval (black bar) displaying within density plot (orange), which depicts the mean difference in CT between groups for each iteration. None of the sulci significantly differed between DPs and NTs ($F(1,468)<2.0$, $p>.15$).

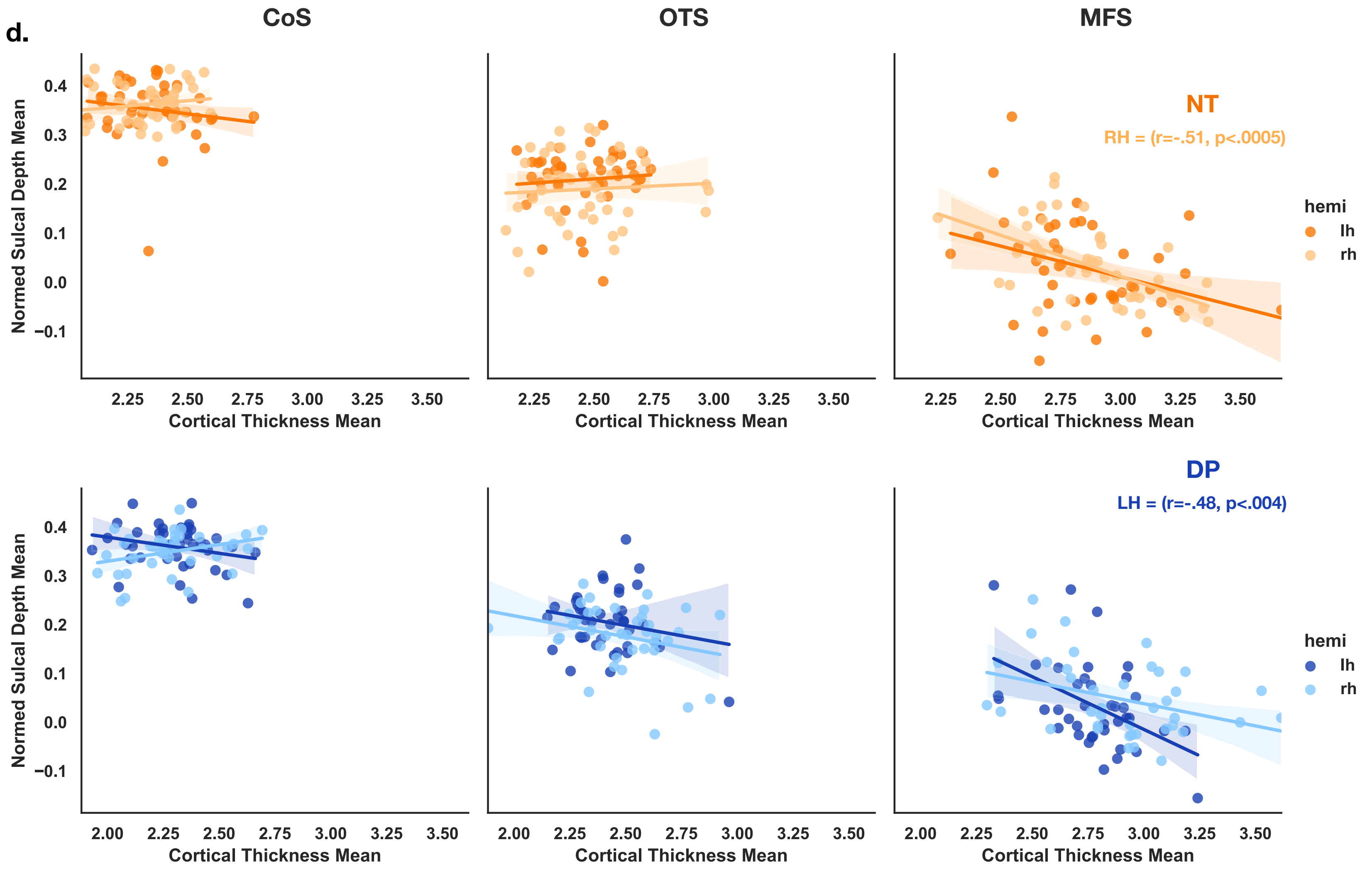

**Supplementary Figure 5. Relationship between cortical thickness and sulcal depth in VTC in DPs and NTs.** Correlations between cortical thickness and sulcal depth for three VTC sulci (CoS, OTS, MFS) are plotted with NTs in the top row (orange) and DPs in the bottom row (blue; LH: darker shade; RH: lighter shade). Only the MFS (right) showed a significant anti-correlated relationship between cortical thickness and sulcal depth in the NT right hemisphere ($r=-.51$, $p<.0005$) and DP left hemisphere ($r=-.48$, $p<.004$).

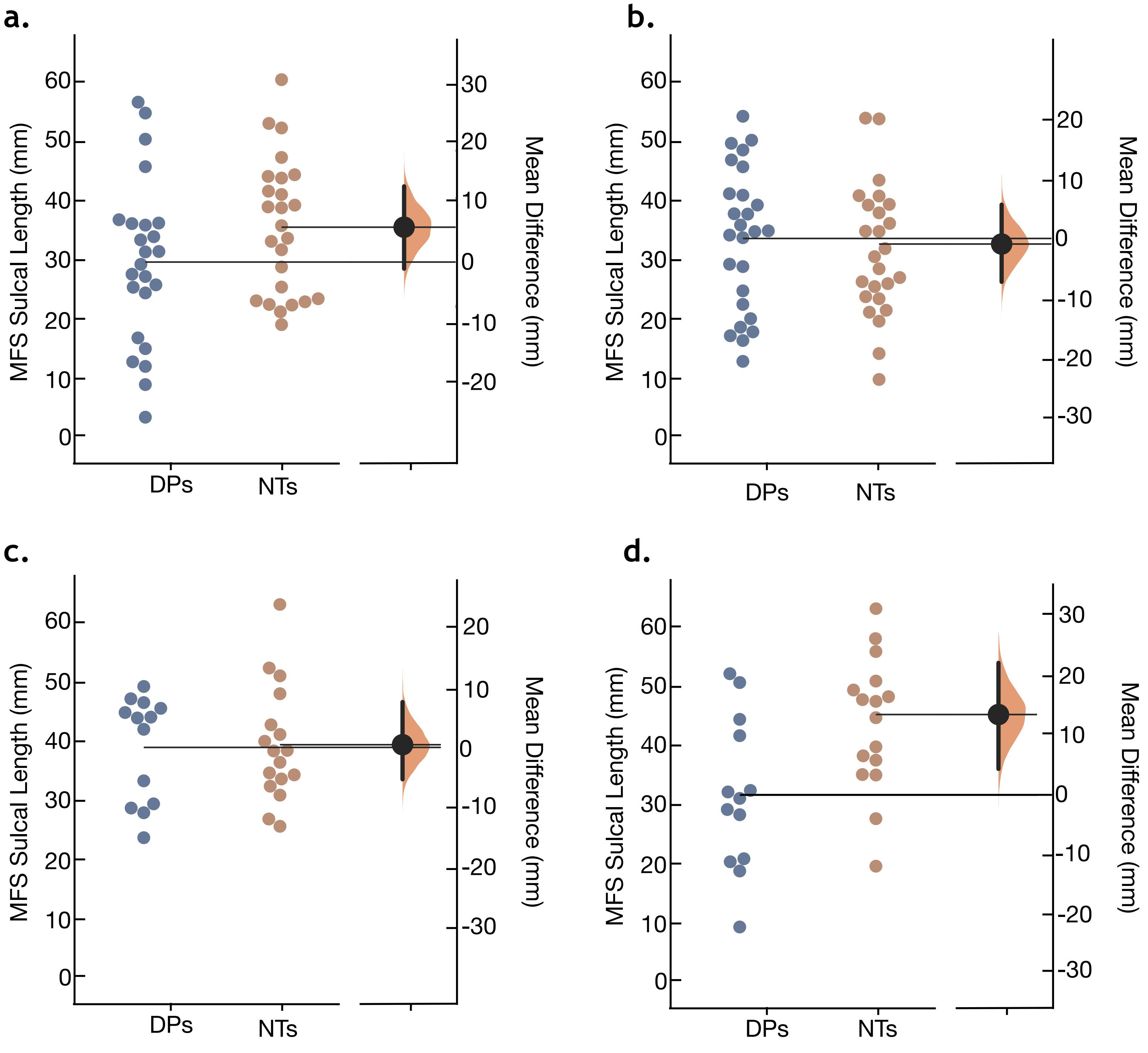

**Supplementary Figure 6. Statistically unpacking a group x hemisphere x gender interaction.** Swarm plot showing MFS sulcal length as a function of group (DPs, blue; NTs, orange), broken down by hemisphere and participant gender (**a.** LH Females; **b.** RH Females; **c.** LH Males; **d.** RH Males). On the right y-axis, a bootstrap of 5,000 iterations was used to generate a confidence interval (black bar) displayed within a density plot (orange), which depicts the mean difference in MFS length for each iteration. The difference in length between DPs and NTs is driven primarily by the male right hemisphere (t=2.9, $p$=.009) and a marginal effect in the female left hemisphere (t=-1.6, p=.11).

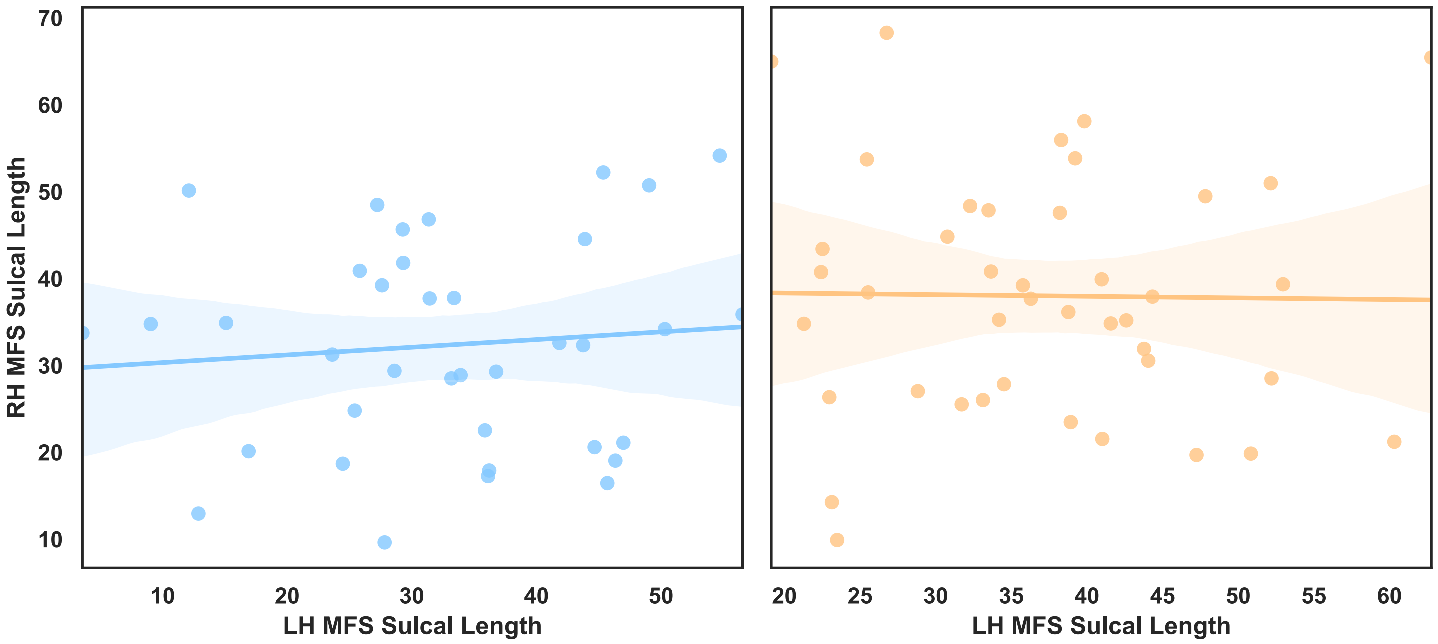

**Supplementary Figure 7. No observed correlation of MFS sulcal length in left and right hemisphere in individual participants.** The length of the MFS in the left hemisphere (x-axis) and right hemisphere (y-axis) hemispheres are plotted in both DPs (blue) and NTs (orange). Neither DPs nor NTs showed a significant relationship between left and right hemisphere MFS length (DP: r=0.1, *p*=0.5; NT: r=-0.01, *p*=0.9).
